# Sperrfy the brain: A data-driven realization of Sperry’s Chemoaffinity theory in the neural connectome

**DOI:** 10.1101/2025.04.30.651442

**Authors:** Jigen Koike, Yuichiro Yada, Riichiro Hira, Honda Naoki

## Abstract

Understanding how brain-wide neural circuits are genetically wired remains a fundamental question in neuroscience. While Sperry’s chemoaffinity theory posits that molecular gradients provide positional cues for axonal projections, its application has been largely limited to localized sensory systems. Here, we present SPERRFY (Spatial Positional Encoding for Reconstructing Rules of axonal Fiber connectivitY), a data-driven framework that operationalizes Sperry’s theory at the whole-brain scale. By integrating connectomic data with spatial transcriptomic profiles from the Allen Mouse Brain Atlas, SPERRFY deciphers latent positional gradients that underlie axonal wiring. Using canonical correlation analysis (CCA), we extract top gradient pairs that align with observed neural connectivity patterns, capturing both global (inter-regional) and local (intra-regional) organizational principles. Connectivity reconstruction based on these gradients achieves high predictive performance (AUC = 0.90), and permutation-based null models confirm the biological relevance of the inferred structures. Furthermore, SPERRFY identifies candidate genes that may contribute to positional wiring information, providing molecular insight into the developmental logic of brain-wide circuitry. Our results extend Sperry’s foundational theory beyond the sensory domain, offering a unified, data-driven framework for understanding genetically encoded connectivity across the entire brain.

## Introduction

The structure of brain circuits is extremely complex and is the basis of higher brain functions such as recognition. One of the most fundamental questions in brain science is how these complex circuits are genetically designed and wired. Many molecular biologists have extensively investigated how neural circuits are wired through axon guidance, in which axons elongate and chemotactically migrate in an attractive or repulsive manner depending on the type of guidance cues^1–3^. So far, a large number of guidance cues, e.g., netrin^4,5^, semaphorin^6^, ephrin^7–13^, and their receptors have been identified. However, these microscopic molecular findings are insufficient for understanding the wiring of the entire brain.

In neural circuit formation, neurons must not only project axons chemotactically, but also correctly recognize their projection destination. In 1963, Roger Sperry proposed the chemoaffinity theory, which states that neurons rely on chemical gradients as positional information to recognize the destination of axon projections^14,15^. Numerous studies supported this chemoaffinity theory and identified a large number of molecules that carry positional information^10^. For example, in the topographic circuit of the visual system, Eph receptors and ephrin ligands are expressed in gradients in the retina (source region) and the tectum (target region), respectively^16–19^ (**Fig. 1a**). The neurons in the source region project their axons to the target region, where they reach the position with a specific ephrin concentration depending on the expression level of Eph receptors in the source region^18,19^. In other words, the concentration gradients of Eph and ephrin are responsible for the “*wiring positional information* (wiring PI)” underlying the genetically programmed neural wiring. However, studies of the chemoaffinity theory have thus far been limited to simple circuits, mainly sensory circuits with topographic structure, e.g., visual and olfactory systems^20,21^. Due to the complexity of brain development, few studies have experimentally investigated complex neural circuits such as those of the cerebrum.

**Figure 1:**
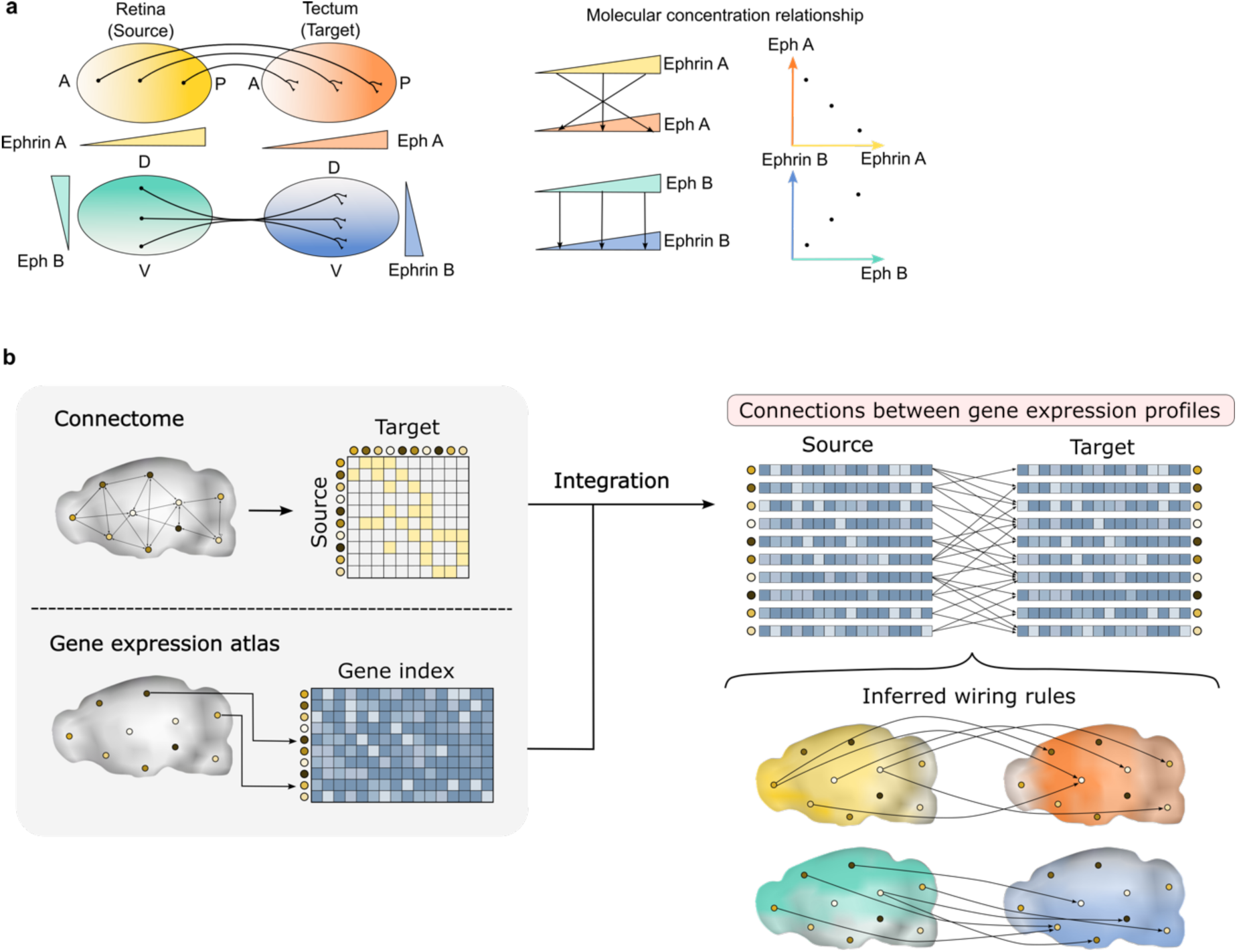
Schematic overview of the chemoaffinity theory and the framework of SPERRFY. **a:** Conceptual illustration of the chemoaffinity theory in the topographic organization of the visual system. (Left) Eph receptors and ephrin ligands are expressed as gradients in the source region (retina) and the target region (tectum), respectively. Axonal projection sites are determined based on the concentration levels of these molecules. Different combinations of molecular gradients along the anterior-posterior (AP) and dorsal-ventral (DV) axes regulate projection specificity. (Right) Schematic representation of the molecular concentration relationship between source and target sites. **b:** Overview of the SPERRFY framework. (Left) SPERRFY integrates connectome data and spatial gene expression data (transcriptome atlas). (Right) By combining both datasets, latent molecular gradients underlying neural wiring can be inferred, revealing projection rules consistent with the chemoaffinity theory.

Meanwhile, the structure of the whole brain has been thoroughly mapped in recent years. The brain’s neural wiring pattern (connectome) has been revealed in several animal models, including humans^22,23^, mice^24^, marmosets^25,26^ and Drosophila^27–30^. In addition, spatial transcriptomic data —the spatial distribution of genome-wide gene expression in the brain— has also been obtained^31–37^. These data on connectomes and spatial transcriptomes are now available in public databases. Many data science studies have investigated such data, revealing the relationship between neural connectome patterns and gene expression^38–41^. However, despite the availability of such data, no studies have focused on the wiring PI of neural circuits from the perspective of chemoaffinity theory.

In this study, we developed a new data-driven framework named “SPERRFY” (Spatial Positional Encoding for Reconstructing Rules of axonal Fiber connectivitY) to decipher wiring positional information in actual brain data based on the chemoaffinity theory (**Fig. 1b**). This method employs a machine learning technique called canonical correlation analysis (CCA)^42^ to extract hidden relationships between gene expression vectors at the source and target of neural projections. By applying our method to data from the Allen Brain Atlas^24,32,33,43^, we extracted pairs of concentration gradients that best explain the actual neural wiring patterns in the mouse brain. These patterns are explained in the context of the chemoaffinity theory as wiring positional information. We then verified the validity of the chemoaffinity theory throughout the whole mouse brain, by reconstructing neural wiring patterns using the identified wiring positional information. Furthermore, we compared these results to those of null models, suggesting that both local and global control mechanisms underlie the brain’s neural circuitry. Therefore, the SPERRFY framework extends the understanding of the chemoaffinity theory from simple sensory circuits to more complex brain regions.

## Results

### Heart of SPERRFY framework

“SPERRFY” (Spatial Positional Encoding for Reconstructing Rules of axonal Fiber connectivitY) is designed to discover wiring positional information (wiring PI), i.e., the latent gradients that best explain actual neural circuits based on the chemoaffinity theory, using paired gene expression vectors at source and target sites of neural connections (**Fig. 2a**). According to the chemoaffinity theory, neural wiring is determined by the molecular concentration gradients between source and target sites, resulting in a high correlation between the molecular concentrations at the source and target sites of axonal projections^19^. Leveraging this property, our method inversely identifies the correlated structures underlying axonal projections through canonical correlation analysis (CCA).

**Figure 2:**
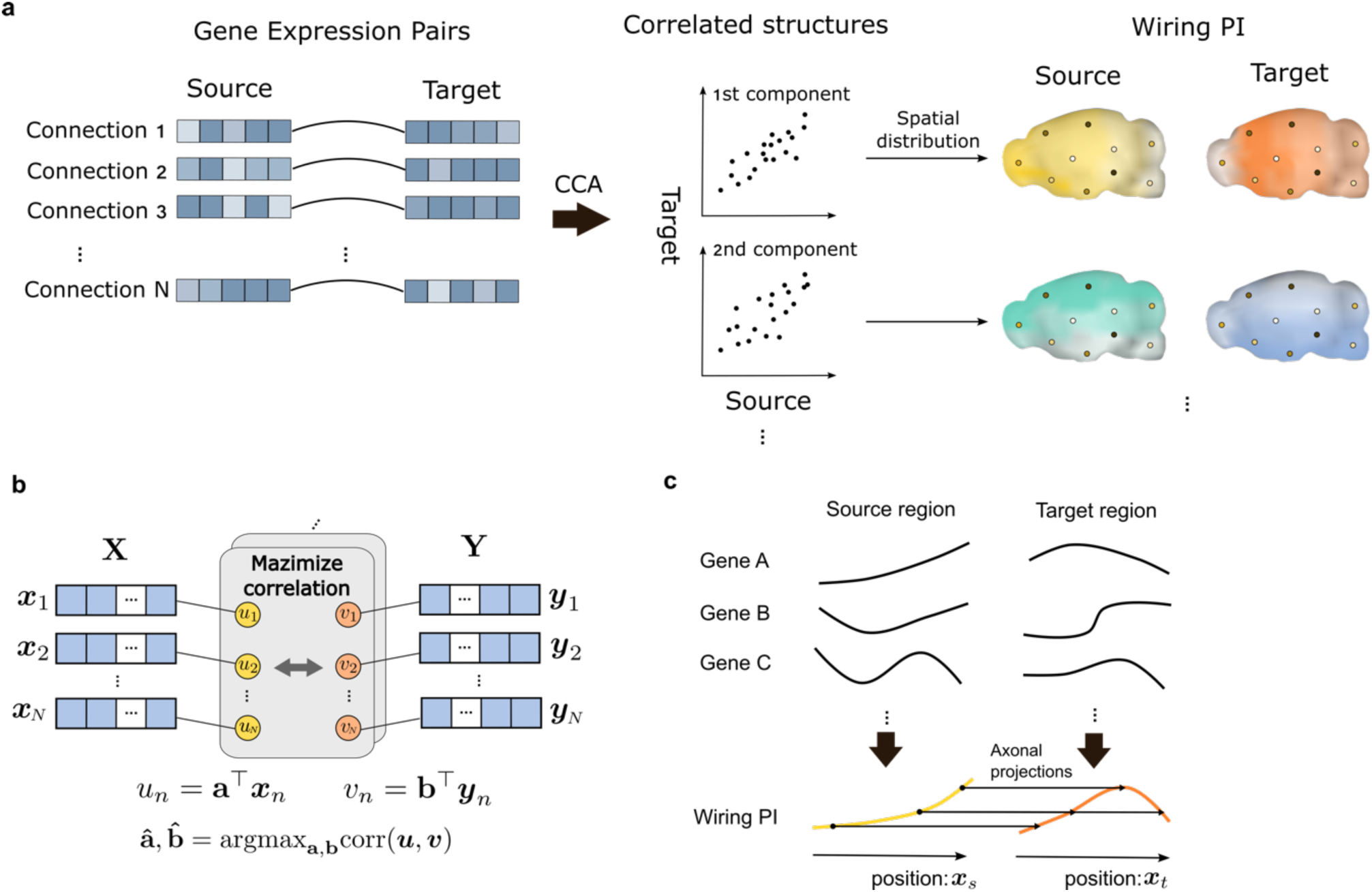
Workflow of SPERRFY and its theoretical basis. **a:** Workflow of the SPERRFY framework. (Left) SPERRFY utilizes integrated connectome and gene expression atlas data, represented as paired gene expression profiles between the source and target sites of neural projections. (Middle) By applying canonical correlation analysis (CCA), SPERRFY identifies multiple correlated structures between the source and target sites. (Right) The spatial distributions of each wiring PI pair are visualized by mapping their gradients onto the brain regions involved in the connections. **b:** Conceptual diagram of canonical correlation analysis (CCA), which identifies pairs of linear combinations from two datasets (**X** and **Y**) that maximize their correlation. **c:** Interpretation of SPERRFY as an extension of the chemoaffinity theory. For simplicity, this model is illustrated in one-dimensional space. In this extended framework, wiring positional information (PI) is assumed to emerge from the combined influence of multiple gene expression patterns, rather than from individual molecular gradients such as Eph/ephrin. Axonal projections are assumed to form between source and target sites with similar PI values.

CCA is an unsupervised machine learning technique for finding correlated structures between paired vector data from different modalities^42^. CCA considers weighted sums of the components of vector data within each modality, and identifies several combinations of weights that achieve the highest correlations between the weighted sums (see Methods, **Fig. 2b**). In general, CCA produces multiple pairs of weights for which the correlation is maximized.

The SPERRFY extends the interpretation of a mathematical model of chemoaffinity theory^19^ by assuming that wiring PI arises from the combinatorial effects of multiple gene expressions (**Fig. 2c**). To align with the CCA framework, the wiring PI is formulated as multiple pairs of the weighted sums of the gene expression data of source (***x***_*s*_) and target (***x***_*t*_) sites of axonal connection:

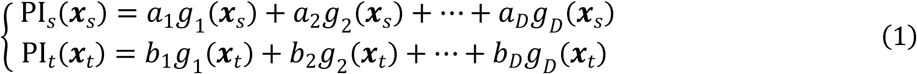

where ***x***_*s*_ and ***x***_*t*_ represent spatial coordinates of source and target brain region, and *g*_*d*_(***x***) denotes the expression levels of the *d* th gene at position ***x***. ***a*** = (*a*_1_, …, *a*_*D*_)^T^ and ***b*** = (*b*_1_, …, *b*_*D*_)^T^ are the weight vectors to be estimated by CCA. SPERRFY assumes that axonal projections occur where PI values of the source and target regions are similar. SPERRFY maximizes correlations between PI_*s*_(***x***_*s*_^(*n*)^) and PI_*t*_(***x***_*t*_^(*n*)^), where ***x***^(*n*)^ = (***x***_*s*_^(*n*)^, ***x***_*t*_^(*n*)^) represents pairs of brain coordinate indices of the *n* th connection (**Fig. 2a**). CCA calculates multiple pairs of 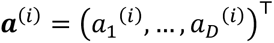 and 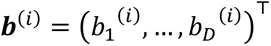 for the *i* th correlated structure, and we can obtain their corresponding gradients of PI_*s*_^(*i*)^(***x***_*s*_) and PI_*t*_^(*i*)^(***x***_*t*_). The wiring PI obtained by SPERRFY reflects the connection patterns in the original connectome data. Furthermore, by reconstructing brain wiring from these positional gradients, we can evaluate the explanatory power of the chemoaffinity theory in actual neural circuits.

### Analysis of mouse brain data from Allen Brain Atlas

For the application of SPERRFY, we used the Allen Brain Atlas, a public database containing neural connectivity data^24^ and gene expression data of adult mice^32,33^. These data can be easily integrated because they have been registered in the same standard brain atlas^43,44^. We transformed these data into paired vector data of gene expression levels to apply the SPERRFY framework (**Fig. 2a**).

The neural connectivity data we used are mesoscale connectome data from work by Oh et al.^24^, which take the form of a binary matrix representing directed connections between 213 brain regions of the right hemisphere (**Fig. 3a**). These 213 regions are further divided into 13 major regions (MRs): isocortex, olfactory areas, hippocampal formation, cortical subplate, striatum, pallidum, thalamus, hypothalamus, midbrain, pons, medulla, cerebellar cortex, and cerebellar nuclei.

**Figure 3:**
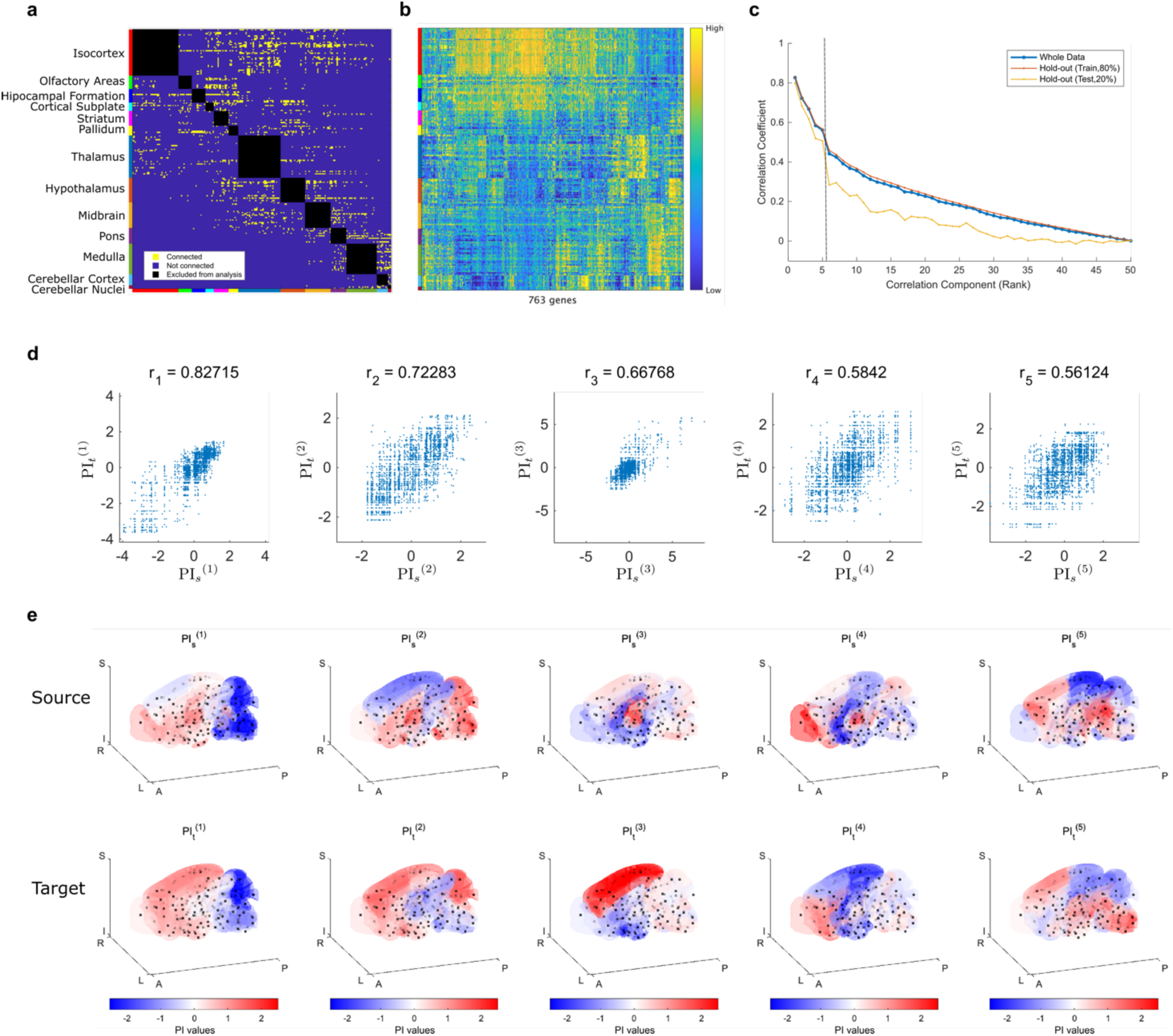
Correlated structures underlying mouse brain wiring extracted by SPERRFY. **a:** Binary directed matrix representing mesoscale neural connectivity in the mouse brain. Connections within the same major region (MR) are excluded from the analysis. **b:** Gene expression matrix showing normalized expression values for 763 genes across 213 brain regions. **c:** Correlation coefficients of the wiring PI components extracted via CCA. Values are ranked in descending order. The blue line indicates results from the full dataset, while the red and yellow lines show results from the hold-out validation (training and test sets, respectively). **d:** Scatter plots of the top five most strongly correlated wiring PI components. Each point represents a single neural connection, with x- and y-values corresponding to PI values at the source and target regions. **e:** Spatial distributions of the top five wiring PI gradient pairs mapped onto the standard mouse brain. Source (top row) and target (bottom row) distributions are shown for each component. Red and blue indicate high and low PI values, respectively.

The gene expression data are a matrix of the expression levels of 763 genes in these 213 brain regions (**Fig. 3b**). In this analysis, gene expression levels were reduced to 50 dimensions using PCA^45^ to summarize genes showing redundant expression distribution (**Supplementary Fig. 1**). We integrated this gene expression data and connectivity data into paired vectors of dimensionally reduced gene expression levels in the source and target region of each connection for analysis.

### Wiring positional information (PI) of entire mouse brain

First, we calculated the wiring positional information of the entire mouse brain connectivity using data from the Allen Brain Atlas^24,32,33,43^. To exclude the effect of gene expression correlations in neighboring regions, we focused our analysis exclusively on connections between different MRs, resulting in a total of 2,213 connections (**Fig. 3a**). In addition to analyzing the entire dataset of 2,213 connections, we performed hold-out validation^46^ by randomly dividing the connection matrix into training and test regions, and computing the correlation structures based solely on the training dataset (See Methods and **Supplementary Fig. 2**).

By applying SPERRFY, we identified latent correlated structures between the source-target pairs of connections, among which the top 5 exhibited particularly strong correlations (**Fig. 3c, d**). We also confirmed that these top 5 correlated structures were consistently preserved in hold-out validation datasets (**Fig. 3c**). Therefore, the following analysis focused on these top 5 correlated structures.

For these correlated structures, we visualized the identified gradients PI_*s*_^(*i*)^(***x***) and PI_*t*_^(*i*)^(***x***) in the standard mouse brain^44^ (**Fig. 3e**). These gradient pairs represent wiring PI and provide the best explanation of the connectome data in the context of the chemoaffinity theory. The most correlated gradient pair exhibits similar distributions, with higher values in the anterior part of the brain and lower values in the posterior part (**Fig. 3e, Supplementary Fig. 3**). The brain’s neural connection matrix exhibits a modular organization, which can be roughly separated into anterior (isocortex, olfactory areas, hippocampal formation, cortical subplate, striatum, pallidum, thalamus, hypothalamus, and midbrain) and posterior (pons, medulla, cerebellar cortex, and cerebellar nuclei) parts (**Supplementary Fig. 3**). This most correlated gradient pair may be associated with this modular organization. The second most correlated pair of gradients differs from the most correlated pair, with significantly different distributions between the source and target gradients (**Fig. 3e**). Notably, some MR pairs— such as isocortex to thalamus, isocortex to midbrain, and midbrain to thalamus—show relatively similar values between source and target, suggesting that this wiring PI may encode preferred directional relationships between specific brain region pairs, rather than reflecting global modularity alone (**Supplementary Fig. 3**). The third most correlated pair exhibits a characteristic distribution, with high variance only in the thalamus for the source and the isocortex for the target (**Supplementary Fig. 3**). This distribution pattern suggests a potential role in controlling projections from the thalamus to the isocortex. The fourth and fifth gradient pairs exhibit greater variance within each MR compared to the top three correlated pairs. These are more likely to reflect finer-scale connection patterns, i.e., intra-MR connections, rather than inter-MR projection density.

### Genes showing similar distribution to wiring PI

To identify genes associated with the wiring PI pairs, we screened for genes whose spatial expression patterns resemble the identified wiring PI gradients. We calculated cosine similarity scores (see Methods) between the spatial expression patterns of 763 genes and each of the extracted wiring PI components (**Fig. 4a**). For each wiring PI, we identified the top-ranked genes and visualized their expression profiles across brain regions (**Fig. 4b**). These genes exhibit distributions similar to the wiring PIs, either in their original form or with inverted signs. These visualized examples illustrate how certain gene expression patterns closely align with the spatial structure of the inferred wiring PIs. The complete rankings of the top 10 genes with the highest similarity scores for each wiring PI gradient are listed in **Table 1**. Note that the rankings are based on the absolute values of the cosine similarity scores, as the sign of the wiring PI values may be flipped without affecting the strength of correlation.

**Figure 4:**
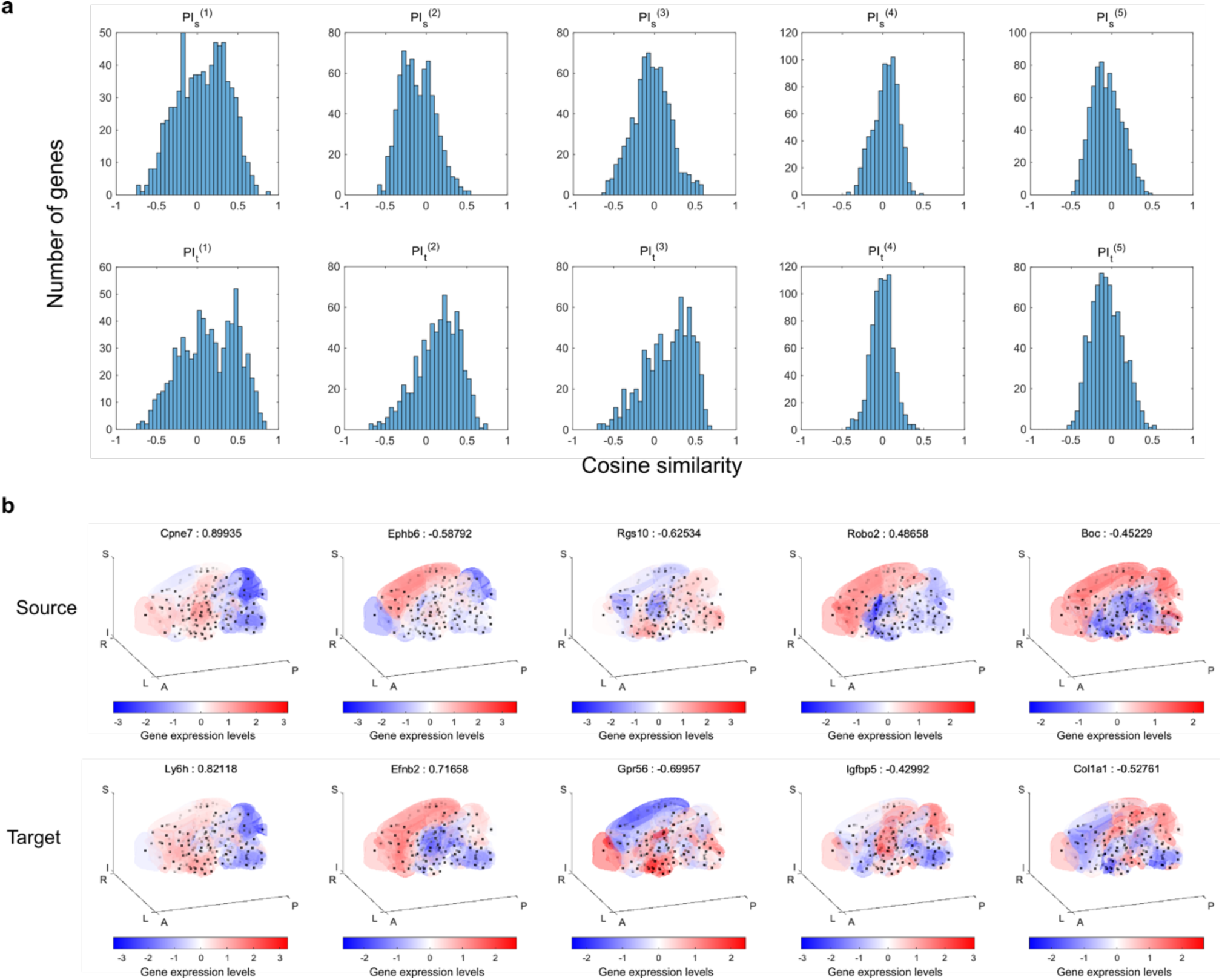
Gene-level correspondence and spatial expression profiles associated with wiring PI gradients. **a:** Histograms showing the distribution of cosine similarity scores between each wiring PI component and the expression patterns of 763 individual genes. The top row corresponds to source PI components (PI_!_), and the bottom row to target PI components (PI_“_). **b:** Spatial gene expression maps of the genes with the highest absolute cosine similarity to each wiring PI gradient. For each component, the most similar gene expression pattern is shown for both source (top) and target (bottom) regions, with the corresponding gene name and similarity score indicated above each map. Red and blue indicate high and low expression levels, respectively.

**Table 1:**
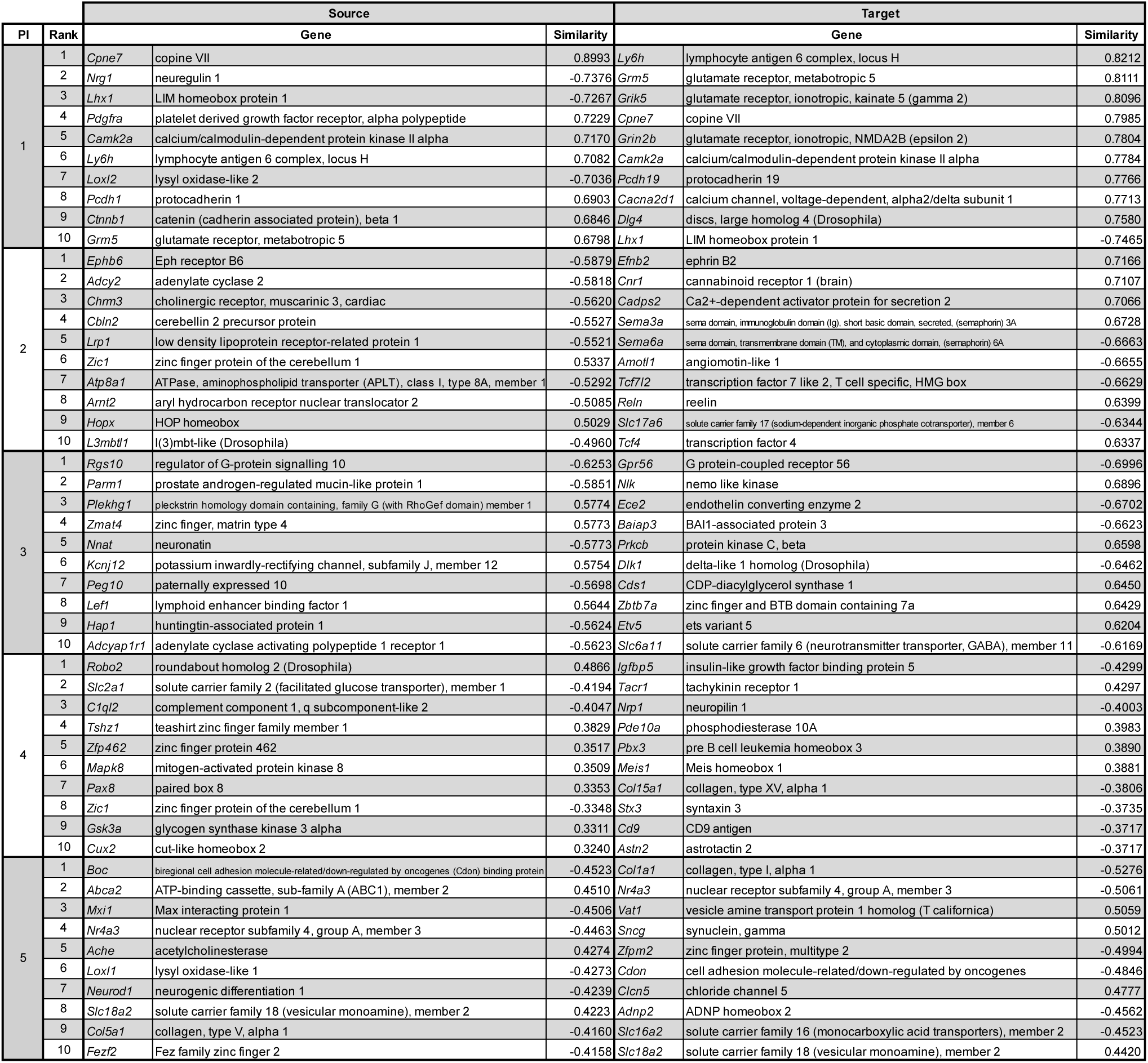
Genes with spatial expression patterns most similar to the identified wiring PI gradients. The table lists the top 10 genes with the highest cosine similarity to each of the top five wiring PI components, separately for source and target regions. Similarity values were computed across 213 brain regions. A higher absolute value of similarity indicates a stronger correspondence between the gene expression pattern and the wiring PI gradient.

Genes with such spatial patterns may play a significant role in guiding axonal projections during brain development by providing positional molecular cues. Notably, for the second PI component, we observed strong similarity to Ephb6 and Efnb2, both members of the Eph/ephrin signaling family known to mediate topographic projections in visual and other sensory systems^10,14,18,19,47^. This supports the interpretation of wiring PIs as molecular gradients consistent with the chemoaffinity theory.

Other components also exhibited correspondence with genes potentially relevant to neural wiring. The first PI component showed strong similarity to genes involved in excitatory synaptic transmission, including *Grm5* (mGluR5)^48,49^, *Grin2b*^50^, *Dlg4* (PSD-95)^51^, and *Camk2a*^52^, as well as genes associated with neural development such as *Nrg1*^53^ and *Pcdh19*^54^. Notably, the source and target PI gradients for this component exhibited highly similar spatial distributions, and several genes—*Camk2a*, *Grm5*, and *Cpne7*—appeared in both rankings. The third PI component showed similarity to *Gpr56*^55,56^, a member of the adhesion-GPCR family known to play essential roles in cortical development and neuronal positioning. The fourth component showed strong similarity to *Robo2*^57^, a key axon guidance receptor that mediates Slit signaling and determines axonal projection trajectories during brain development. The fifth component included *Boc*^58,59^, a co-receptor for Sonic Hedgehog (Shh) signaling involved in commissural axon guidance. Together, these results demonstrate that the spatial structure of wiring PIs recapitulates the distribution of genes known or suspected to participate in neural circuit formation, suggesting that SPERRFY captures biologically meaningful positional information.

### Reconstructing connectome structure using wiring positional information

Next, to verify the chemoaffinity theory in the actual whole-brain circuits of mouse, we tested whether the identified wiring PI, derived from gene expression data, could be used to reconstruct the neural connectome structure. In the classical chemoaffinity theory, the location of axonal projections is determined by specific molecular concentration gradients (e.g., Eph receptors and ephrin ligands) at the source and target regions. Our SPERRFY framework identifies such concentration gradient pairs as wiring PI from the brain data with the help of CCA. However, it remains uncertain how well the identified wiring PI gradients reflect the actual wiring patterns of the connectome data. To address this issue, we reconstructed neural wiring patterns from identified wiring PIs and compared them to the original connectome matrix.

In reconstructing the wiring patterns, we made the following assumption: brain region pairs with similar source and target wiring PI values are more likely to have neural connections, whereas those with dissimilar values are less likely. Based on this assumption, we assessed the extent to which every region pair is likely to be connected from the identified wiring PI gradients. Specifically, we calculated the total difference in the top 5 identified wiring PI gradients as *d*_*PI*_(***x***_*s*_, ***x***_*t*_) (See Methods, equation (16)) (**Fig. 5a**). We then reconstructed the binary connection matrix by determining a connection when the value is below the threshold (**Fig. 5b**).

**Figure 5:**
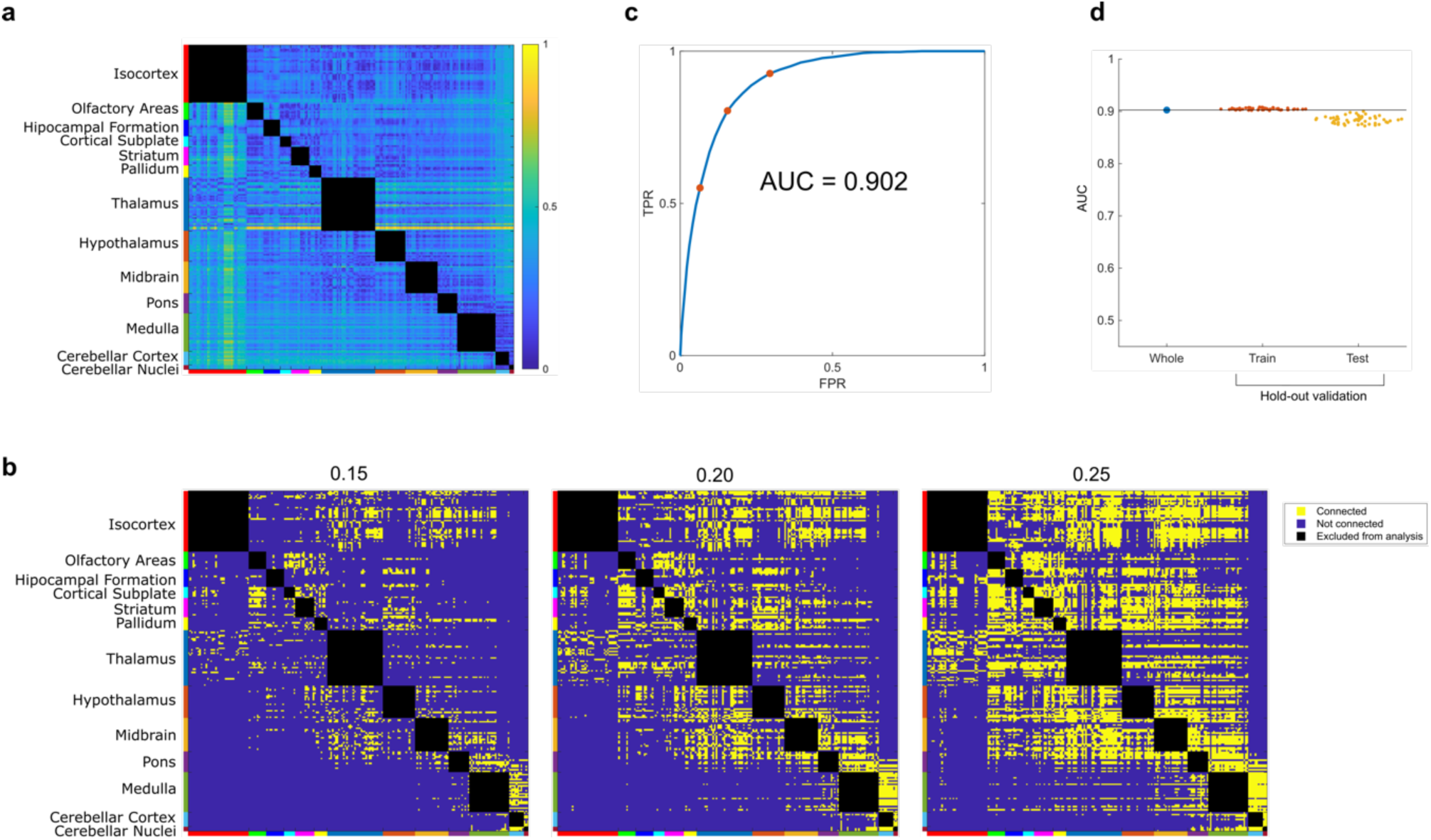
Reconstruction of neural connectivity from wiring PI gradients. **a:** Matrix used for connection reconstruction, representing the squared Euclidean distance between the top five wiring PI gradients for each region pair. Lower values indicate a higher likelihood of neural connectivity. **b:** Examples of reconstructed binary connection matrices using threshold values of 0.15, 0.20, and 0.25, respectively. **c:** Receiver operating characteristic (ROC) curve for connection reconstruction. The area under the curve (AUC) is 0.902. Red dots correspond to the three threshold conditions used in (b). **d:** AUC values for connection reconstruction evaluated on the training and test sets in hold-out validation.

To evaluate the performance of this reconstruction, we generated binary connection matrices with various thresholds and compared them to the original connectome patterns. Then we plotted the receiver operating characteristic (ROC) curves of the connection discrimination by computing the true positive rate (TPR) and false positive rate (FPR). The area under the curve (AUC) of this ROC curve was 0.90, indicating that the connection reconstruction matrix can predict actual connection patterns with high accuracy (**Fig. 5c**). We also confirmed that the prediction performance was robust when applied to the hold-out validation dataset. (**Fig. 5d**). These results indicate that the chemoaffinity theory plays a significant role in explaining the actual wiring structure of the whole mouse brain, and that spatial patterns of gene expression contain information that defines neural wiring structure.

### Comparison of globally and locally randomized neural connection patterns

The above results provide substantial support for the chemoaffinity theory as a model explaining the wiring structure of the mouse brain. Nevertheless, it remains possible that SPERRFY could extract correlated wiring PI gradient structures even from randomly wired connectome data. To evaluate this possibility, we applied SPERRFY to randomized connection patterns. Specifically, we developed a null model, referred to as the “globally randomized model,” which generates globally randomized connection patterns while preserving the original number of connections (**Fig. 6a**, left). Using this model, we generated 1,000 randomized connection patterns and applied SPERRFY to each. We confirmed that SPERRFY could not extract correlated wiring PI structures from these random patterns (**Fig. 6b, c,** red line and dots), resulting in extremely low reconstruction performance (red dots in **Fig. 6d**). These findings demonstrate that the SPERRFY framework does not simply extract highly correlated structures by chance, nor does it achieve successful connection reconstruction for the mouse brain connectivity by coincidence.

**Figure 6:**
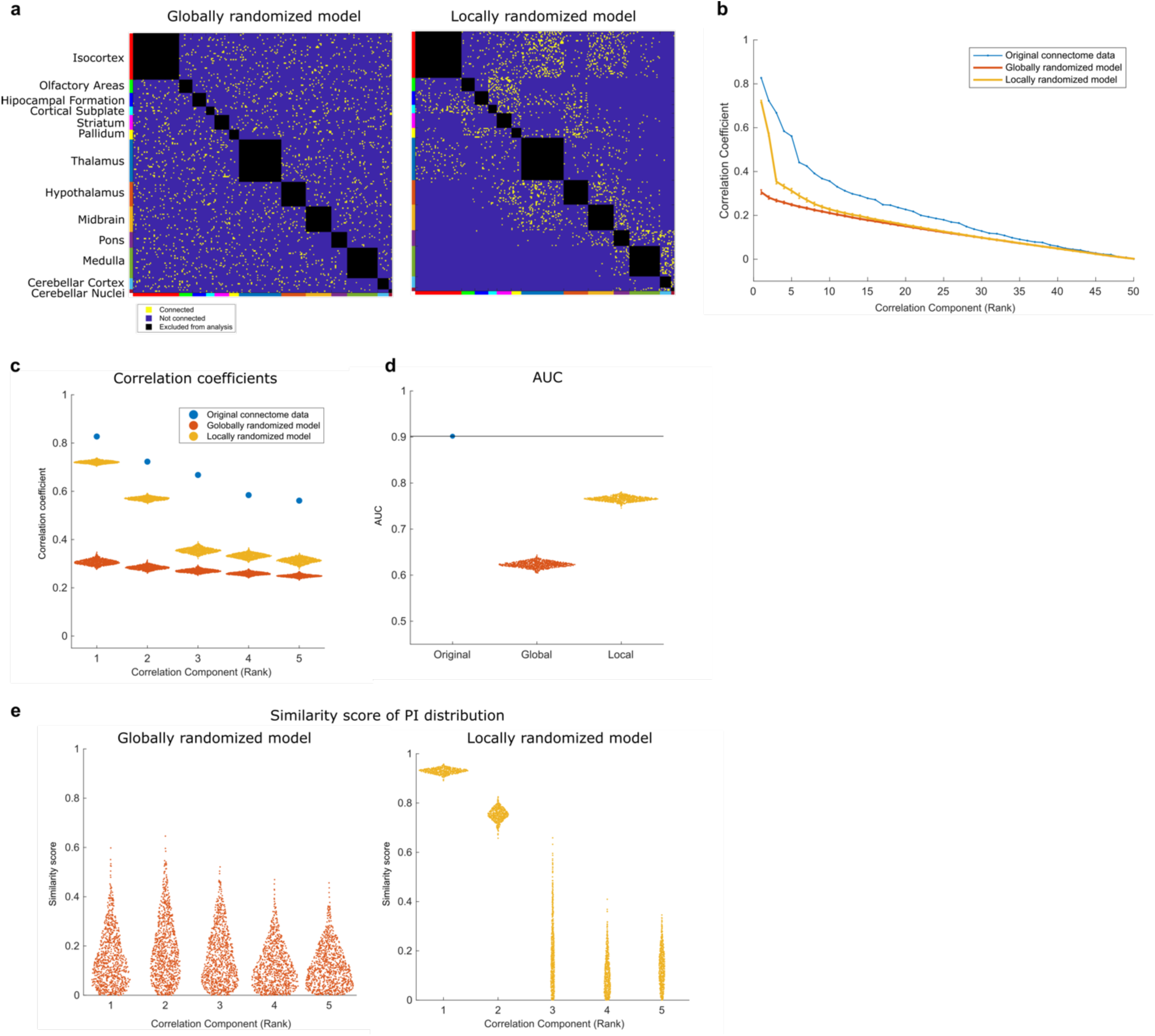
Comparison of SPERRFY results with randomized connection models. **a:** Example binary connection matrices generated from the globally randomized model (left) and the locally randomized model (right). **b:** Mean correlation coefficients of SPERRFY-extracted wiring PI components from the original and randomized connectomes. The blue line represents the original data, the red line the globally randomized model, and the yellow line the locally randomized model. Error bars indicate standard deviations. **c:** Distributions of the top five correlation coefficients extracted from each dataset. Color coding matches that in panel (b). **d:** Distributions of AUC values for reconstructed connectivity across original and randomized models. **e:** Similarity scores of wiring PI gradient distributions between randomized and original data. Left: globally randomized model. Right: locally randomized model.

The globally randomized model served as a useful null model above, but it has limitations in distinguishing global from local effects in neural wiring patterns because it disrupts macroscale connectivity including the brain’s modular organization. We thus developed the “locally randomized model” that disrupts only local connections within each MR pair, while maintaining macroscale connectivity (see Methods). Then we applied SPERRFY to the locally randomized patterns (**Fig. 6a**, right). The correlation between wiring PI gradients and reconstruction performance were both higher than those of the globally randomized model, but did not achieve the scores of the original data (**Fig. 6b-d**, yellow line and dots). Interestingly, we found that the top two correlated structures robustly exhibited high correlations comparable to those observed in the original connection patterns (**Fig. 6c**). Moreover, the wiring PI gradients of these correlated structures exhibited a similar spatial profile in brain tissue to those obtained from the original mouse brain data (**Fig. 6e**, see Methods). These results indicate that the top two wiring PI pairs are not affected so much by local connections within MRs, but reflect global connections between different MR pairs. In contrast, the third to fifth correlated structures of this null model showed significantly lower correlations than those of the original mouse data (**Fig. 6b, c**). This means that the third to fifth wiring PIs of the original mouse brain capture information related to local connections, and their correlation structures are disrupted in the locally randomized model. Taken together, these findings suggest the existence of two types of wiring PI—those representing global and local connection patterns— highlighting their potential roles in regulating neural connectivity at multiple scales.

In the above analysis, the globally and locally randomized model were employed to validate the wiring PI extracted from the original mouse brain data. However, these null models do not account for key properties of the actual mouse brain, such as connection distances and network topology^60^. To evaluate the impact of these factors on the results of SPERRFY, we introduced additional constraints on such properties into the null models (see Methods). The results showed that these additional conditions did not significantly alter the trends in correlated structures, although slightly higher correlations and reconstruction performance were observed (**Supplementary Fig. 4**). These findings confirm that the wiring PI extracted from the mouse brain data captures substantial and unique information intrinsic to the brain’s actual connectivity.

## Discussion

In this study, we elucidated how genetically programmed positional information shapes neural wiring across the entire brain by developing the SPERRFY framework as an extension of Sperry’s chemoaffinity theory. By applying SPERRFY to the mouse brain data from the Allen Brain Atlas, we decoded the wiring PI gradients explaining neural wiring patterns according to the chemoaffinity theory. We also successfully reconstructed the original wiring pattern from the spatial profile of gene expression, thereby validating the SPERRFY framework. In addition, we uncovered a hierarchy in wiring PI gradients regulating neural wiring at both global and local scales. Therefore, we concluded that SPERRFY not only validates chemoaffinity theory in actual brain data but also provides deeper insights into fundamental wiring principles in the brain.

The proposed SPERRFY framework makes three significant contributions to the field of developmental neuroscience. First, it extends the chemoaffinity theory and applies it to the analysis of complex axonal wiring across multiple brain regions, including higher-order cortical areas. Previous studies have primarily focused on relatively simple and topographic circuits, such as projections from the retina to visual areas^16,17^, peripheral inputs to the somatosensory cortex^61–64^, olfactory circuits^65–69^, and thalamocortical projections^12,70–72^. In contrast, complex circuits, such as cortico-cortical networks, have rarely been investigated experimentally due to their complexity. In this study, we elucidated wiring principles in the whole brain, including these intricate circuits, that were previously considered beyond the scope of investigation.

Second, SPERRFY provides opportunities to experimentally explore molecules involved in neural wiring. While many molecules, such as Eph and ephrin, have been identified as key regulators of topographic circuits in sensory systems^7–9,11–13^, few molecules have been identified in complex networks such as the cerebral cortex. The SPERRFY framework facilitates the identification of candidate genes responsible for neural wiring by linking the extracted wiring PI to gene expression patterns. Therefore, this study bridges molecular findings with systems-level brain architecture.

Third, the SPERRFY method can be applied to other species and datasets. Although this study focused on the mouse brain, recent advances have developed public databases of neural connectomes and spatial transcriptomes for other species, such as humans^22,23,31^, Drosophila^27–30,37^, and marmosets^25,26,36^. Additionally, advancements in measurement technologies are yielding higher-resolution datasets. Applying SPERRFY to such datasets could uncover universal wiring principles across species or reveal species-specific characteristics. The SPERRFY method holds promise as a versatile tool for decoding neural wiring rules across species and developmental stages.

Despite extensive research on the relationship between the brain connectome and gene expression^39^, few computational studies have addressed the wiring structure of the connectome within the framework of chemoaffinity theory. While some studies have attempted to predict neural connections based on gene expression profiles ^73,74^ or cell-type composition^75^, these approaches primarily focus on data-driven prediction and do not explain how molecular gradients might shape brain-wide connectivity patterns. In contrast, SPERRFY reconstructs connection patterns by directly modeling positional molecular cues, offering a mechanistic interpretation of wiring principles grounded in classical chemoaffinity theory.

Other studies have investigated the statistical relationships between brain connectivity and gene expression using data science approaches, without aiming to predict or reconstruct connection patterns.^39^. French and Pavlidis (2011) examined the global correspondence between transcriptomic similarity and anatomical connectivity^38^, while Fulcher et al. (2016) identified transcriptional signatures associated with hub-like regions^40^. Unlike these descriptive approaches, SPERRFY extracts latent correlated structures between the source and target regions of neural projections, providing biologically interpretable gradients that explain connectivity patterns and highlight candidate genes potentially involved in axonal wiring.

The use of brain atlas data necessitates careful consideration of spatial autocorrelation in gene expression to avoid biased results^76^. To address this concern, we implemented null models to generate randomized connectome patterns with preserving the actual distribution of neural connection distance. In addition, we implemented other null models designed to account for the network topology of the connectome data because it is also an important characteristic of the actual brain^77,78^. Through statistical testing with these null models, we confirmed significant correlation between the wiring PI gradients extracted by the SPERRFY, indicating that the results are not biased but reliable.

Although this research provides deeper insights into neural circuit formation, several limitations should be acknowledged. First, we used data from adult mice and did not address dynamic changes during development. This limitation may be mitigated in future studies, as transcriptome data from developmental stages are becoming increasingly available^79–83^, although they will need to be matched to adult data for comprehensive analysis. Additionally, this study primarily focused on correlations and did not establish causal relationships. Future research could address this limitation by experimentally validating the candidate genes identified through SPERRFY. Despite these limitations, this study offers a robust foundation for future investigations into the molecular mechanisms underlying neural circuit formation, paving the way for new discoveries in neuroscience.

## Methods

### Mouse connectome data

The mouse connectome data were obtained from the Allen Mouse Brain Connectivity Atlas^24^. These connectome data derive from a series of viral microinjection experiments that visualize axonal projections at the mesoscale level and are summarized as a matrix representing connectivity between brain regions. We analyzed the 213 × 213 sub-matrix for the right hemisphere, as listed in Supplementary Table 3 of Oh et al.^24^ Following earlier analyses of the same dataset (e.g., Ji et al. 2014^73^; Fulcher & Fornito 2016^40^), we binarized the matrix by retaining only those edges with a projection-strength *p*-value < 0.05, yielding 3,123 directed connections.

The 213 brain regions in the connectivity matrix are grouped into 13 major regions (MRs): isocortex, olfactory areas, hippocampal formation, cortical subplate, striatum, pallidum, thalamus, hypothalamus, midbrain, pons, medulla, cerebellar cortex, and cerebellar nuclei. In our analysis, we focused on inter-MR connectivity, excluding intra-MR connections, in order to minimize the effect of correlated gene expression within the same MR. As a result, the inter-MR connectivity matrix includes 2,213 connections.

### Gene expression data of mouse brain

The gene expression data were obtained from the publicly available Allen Mouse Brain Atlas database^32,33^. This database provides summarized results of numerous in situ hybridization (ISH) experiments conducted on the adult mouse brain. Using the Allen API (api.brain-map.org/api/v2/data), we retrieved gene expression datasets corresponding to the same set of 213 regions as used in the connectome data. The details of the data acquisition process are described below.

First, we retrieved identification numbers for all valid ISH experiments (section IDs) by using the following query:

api.brain-map.org/api/v2/data/query.json?

`criteria=model::SectionDataSet,rma::criteria,`

`[failed$eq’false’][expression$eq’true’],products[id$eq1]&num_rows=all`

This query returned 22,157 section datasets containing section IDs. Using these section IDs, gene metadata for each experiment was obtained by querying the following format, replacing “*SectionId*” with the retrieved values:

api.brain-map.org/api/v2/data/query.json?

criteria=model::Gene,rma::criteria,data_sets[id$eq*SectionId*]&num_rows=all

To acquire summarized gene expression data for the 213 brain regions, we retrieved structure IDs for all brain structures in the atlas using the following query:

api.brain-map.org/api/v2/data/query.json?

`criteria=model::Structure,rma::criteria,[graph_id$eq1]&num_rows=all`

This query yielded 1,327 hierarchical brain structures. We then matched the acronyms in the connectivity matrix to identify the structure IDs corresponding to the 213 brain regions. Finally, gene expression levels for these regions were obtained using the following query format:

api.brain-map.org/api/v2/data/query.json?

`criteria=model::StructureUnionize,rma::criteria,`

structure[id$eq*StructureId*]&num_rows=all

where “StructureId” was replaced with the identified structure IDs. Since the retrieved expression data included unsuccessful experiments, we excluded such data by filtering based on the section IDs of the 22,157 section datasets. Additionally, datasets containing missing values for any of the 213 regions were excluded, resulting in a final set of 9,111 genes. To focus our analysis on developmental genes, we further restricted our dataset to genes listed in the Allen Developing Mouse Brain Atlas^35^, ultimately selecting 763 gene expression datasets.

The datasets of gene expression levels include two types of quantified values: expression density and expression energy. In our analysis, we used expression energy, following previous studies^40,74^. Since ISH measures relative rather than absolute expression levels, we normalized the gene expression data using log transformation and z-scoring. To address redundancy in the distribution of the 763 gene expression datasets, principal component analysis (PCA) was applied. PCA was implemented using the `pca` function in MATLAB. For the primary analysis, the top 50 principal components (PCs) were selected based on their contribution rates (**Supplementary Fig. 1**). Additionally, different numbers of PCs (10, 30, 50, 75, and 100) were tested to confirm the effects of the number of PCs (**Supplementary Fig. 5**).

### Integration of connectivity and gene expression data

For the CCA analysis, the connectivity and gene expression data were integrated. Since both datasets shared the same 213 brain regions, this enabled the creation of paired vectors of gene expression levels for all neural connections. Specifically, the binary connectivity data were combined with PCA-reduced gene expression levels from the source and target regions, generating a dataset of paired vectors that served as input for the SPERRFY method. To ensure numerical stability in the CCA computation, a small isotropic Gaussian noise (σ = 0.01) was added to each of the paired gene expression values, breaking near-linear dependencies without altering the resulting wiring-PI gradients.

### Canonical correlation analysis (CCA)

Canonical correlation analysis (CCA) is a machine learning method designed to uncover correlated structures between two sets of variables. Let us consider two multivariate datasets, **X** = (**x**_1_, **x**_2_, …, **x**_*N*_)^T^ ∈ R^*N*×*p*^ and **Y** = (**y**_1_, **y**_2_, …, **y**_*N*_)^T^ ∈ R^*N*×*q*^, where *N* represents the number of paired observations, and *p* and *q* are the numbers of variables (i.e., dimensions) in each dataset. CCA seeks pairs of weighted sums, *u*_*n*_ = ***a***^T^**x**_*n*_ and *v*_*n*_ = ***b***^T^**y**_*n*_, such that the correlation between ***u*** = (*u*_1_, *u*_2_, …, *u*_*N*_)^T^ ∈ R^*N*×1^ and ***v*** = (*v*_1_, *v*_2_, …, *v*_*N*_)^T^ ∈ R^*N*×1^ is maximized. Here, ***a*** ∈ R^*p*×1^ and ***b*** ∈ R^*q*×1^ are weight vectors optimized by CCA. In other words, CCA addresses the following optimization problem:

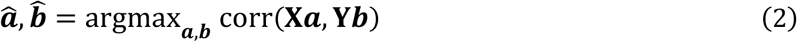

CCA provides not just a single pair of solutions but multiple pairs of 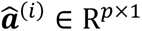 and 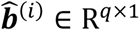, each associated with a correlated pair of 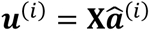 and 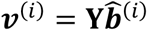. These weight vector pairs are mutually orthogonal (i.e., 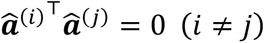), allowing independent patterns of correlation to be extracted. The index (*i*) is ordered according to the magnitude of the corresponding correlation coefficients.

The derivation of the CCA solution is described as follows. For simplicity, **X** and **Y** are assumed to be mean-centered. The correlation coefficient ρ(***a***, ***b***), which is maximized in CCA, is expressed as:

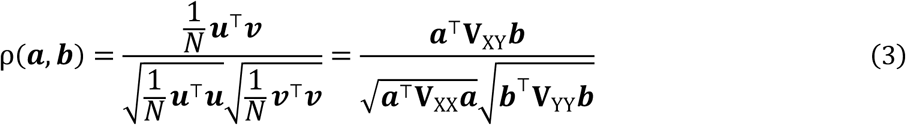

where 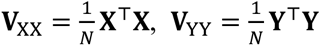, and 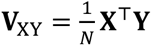 represent the sample variance-covariance matrices of **X** and **Y**, respectively. In CCA, the objective is to find a pair (***a***, ***b***) that maximizes ρ(***a***, ***b***).

Since multiplying ***a*** and ***b*** by positive constants does not change ρ(***a***, ***b***), the standard deviations in the denominator can be normalized to 1. Thus, the optimization problem can be rewritten as:

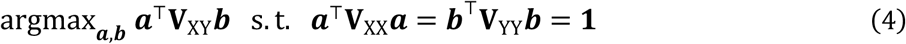

This constrained optimization problem can be reformulated using the method of Lagrange’s multipliers as follows:

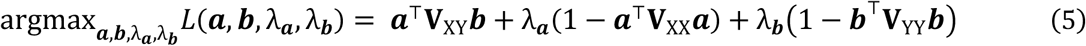

Taking the partial derivatives of *L*(***a***, ***b***, λ_***a***_, λ_***b***_) with respect to a and b and setting them to zero yields:

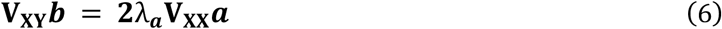

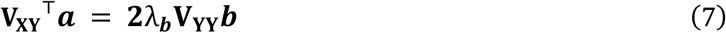

By multiplying both sides of the equations by ***a***^T^and ***b***^T^, respectively, we obtain the following relationship:

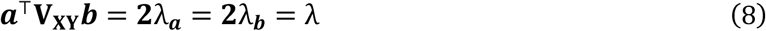

Thus, the above equations can be expressed together in matrix form as:

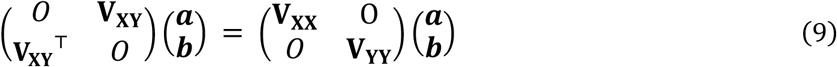

This is a generalized eigenvalue problem, where the eigenvalue λ corresponds to the correlation coefficient ***a***^T^**V**_**XY**_***b***. By focusing on the second and subsequent eigenvalues, up to min{*p*, *q*} number of correlated components can be obtained.

### Application of CCA in the SPERRFY framework

In the framework of SPERRFY, CCA is employed to identify latent correlations between gene expression profiles at the source and target regions of neural connections. Given the paired gene expression data from the source and target regions, CCA extracts linear combinations that maximize their correlation, revealing underlying patterns of wiring positional information (PI).

Let ***x***_*s*_^(*n*)^ and ***x***_*t*_^(*n*)^ represent the spatial coordinate indices of the source and target regions for the *n*th neural connection, respectively. The gene expression levels at these coordinate indices are denoted as ***g***(***x***_*s*_^(*n*)^) ∈ R^*D*×1^ and ***g***(***x***_*t*_^(*n*)^) ∈ R^*D*×1^, where *D* is the number of dimensions after dimensionality reduction. In SPERRFY, the wiring PI is represented as pairs of weighted sums of gene expression levels at the source and target regions:

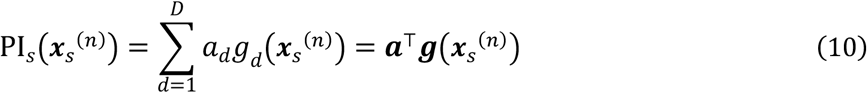

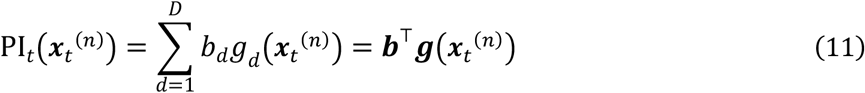

CCA estimates the weight vectors ***a*** ∈ R^*D*×1^ and ***b*** ∈ R^*D*×1^ such that the correlation between the wiring PI at the source and target regions is maximized as

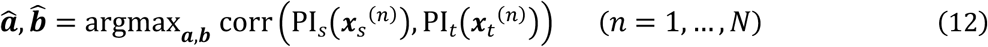

The CCA framework yields multiple pairs of weight vectors 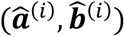, each corresponding to the *i*th highest correlation. With these weights, multiple PI gradients across the brain can be computed as:

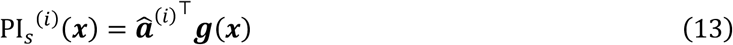

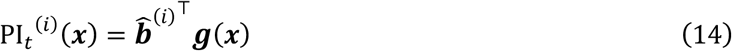

where ***x*** indicates a spatial coordinate of the brain. CCA was implemented using the `canoncorr` function in MATLAB.

### Hold-out validation with train-test split dataset

To assess the reliability of the wiring PI components extracted by SPERRFY, we conducted hold-out validation (**Supplementary Fig. 2**). For this validation, the entire neural connectivity matrix was randomly split into a training set (80%) and a test set (20%). In the training phase, CCA was applied exclusively to the paired gene expression data corresponding to the neural connections in the training set, excluding connections in the test set. This ensured that the extracted PI gradients were derived solely from the training data, without contamination from the test set connections. The extracted wiring PI gradients from the training set were then evaluated on the test set by calculating the correlation coefficients between the PI values at the source and target regions of the test set connections (**Supplementary Fig. 2**). Hold-out validation was also conducted to evaluate the reconstruction of neural wiring from the estimated wiring PI gradients (**Fig. 5d**). This analysis provided a quantitative measure of how well the identified PI gradients capture positional relationship in neural connectivity patterns within unseen data.

### Similarity between gene expression patterns and wiring PI gradients

To screen genes associated with wiring PI, we evaluated the similarity between gene expression patterns and wiring PI gradients using cosine similarity. Given two *D*-dimensional vectors ***x*** and ***y***, cosine similarity *S*_*c*_(***x***, ***y***) is defined as follows:

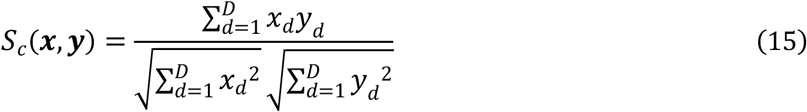

where *x*_*d*_ and *y*_*d*_ are the *d*th components of vectors ***x*** and ***y***, respectively.

For each wiring PI gradient, we computed the cosine similarity between its spatial distribution and the expression pattern of each gene across the 213 brain regions. Similarity scores were evaluated using the absolute value of cosine similarity to assess the strength of spatial correspondence, while the sign of the original score was retained to distinguish between positive and negative associations.

### Reconstruction of neural connection matrix

To evaluate the validity of the chemoaffinity theory in explaining neural circuit formation, we reconstructed the neural connectivity matrix using the wiring PI gradients identified by the SPERRFY framework. According to the chemoaffinity theory, the positions of axonal projections are determined by specific molecular concentration mappings between the source and target regions. In the framework of SPERRFY, these molecular mappings are represented by the proximity of the wiring PI values at the source and target regions. Notably, since the wiring PI gradients obtained through CCA are standardized and expressed in terms of positive correlation, comparing their magnitudes provides a reasonable measure of positional relationships.

First, we quantified the differences in the identified wiring PI values for each pair of brain regions. For the top five most correlated PI components PI_*s*_^(*i*)^(***x***_*s*_) and PI_*t*_^(*i*)^(***x***_*t*_), the L2 norm of their absolute differences was calculated as:

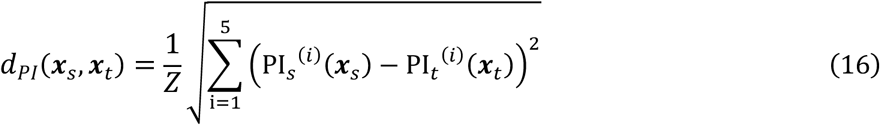

where ***x***_*s*_ and ***x***_*t*_ represent the spatial coordinates of the source and target brain regions, respectively, and *Z* is the normalization term. *d*_*PI*_(***x***_*s*_, ***x***_*t*_) is normalized such that its maximum value is set to 1, ensuring a consistent range for further analysis. A value closer to 0 in this matrix indicates a higher likelihood of neural connectivity.

Second, we reconstructed connectivity matrices by binarizing the *d*_*PI*_(***x***_*s*_, ***x***_*t*_) matrix with a threshold, where region pairs with PI differences below the threshold were classified as connected. To generate the full receiver operating characteristic (ROC) curve, the decision threshold was systematically swept from 0 to 1 in 0.01 increments (101 operating points). The accuracy of the reconstructed connectivity was evaluated by comparing these thresholded matrices to the original connectome. The performance was assessed using the area under the curve (AUC), providing a quantitative measure of how well the reconstructed matrices predict actual neural connections.

### Null models for evaluating wiring positional information

To verify whether the wiring PI identified by SPERRFY reflects biologically meaningful connectivity patterns rather than occurring by chance, we performed statistical testing by employing two types of null models: the globally randomized model and the locally randomized model.

In the globally randomized model, connections were randomly reassigned across the entire connectivity matrix (without any structural constraints), while preserving the total number of neural connections in the original connectome (**Fig. 6a, left**). This approach generated connectivity matrices where the connections were uniformly distributed across all brain regions. In contrast, the locally randomized model imposed structural constraints by preserving not only the total number of neural connections but also the number of connections between MRs (**Fig. 6a, right**). Specifically, connections were randomly reassigned within each MR pair while maintaining the original connection counts for each pair, thereby retaining macroscale connectivity patterns such as inter-region projection density.

To further refine the analysis, two additional constraints were individually applied to the null models in order to better reflect key features of the mouse brain connectome. First, a constraint was introduced to preserve the connection distance distribution by utilizing data from Oh et al., 2014^24^, which provides detailed information on the spatial organization of neural connections in the mouse brain. Connection distances were binned into histograms with a bin width of 100 μm, and connections were randomly reassigned while maintaining the original distribution across these bins. Second, another constraint was applied to preserve network topology by generating randomized connectivity matrices through shuffling the order of brain region annotations while maintaining the original connection patterns. This procedure retained important topological features of the network, such as degree distribution and hub structures, while randomizing the association between connectivity patterns and gene expression profiles.

For each null model, 1,000 randomized connectivity matrices were generated, and SPERRFY was applied to extract wiring PI gradients from each instance. Subsequently, the empirical null distribution of the correlation coefficients between the extracted PI gradients and the accuracy of the reconstructed connectivity was obtained. Statistical significance was assessed by comparing these null distributions with those derived from the original connectome data.

### Evaluation of the similarity of wiring PI gradients

To evaluate the similarity between wiring PI gradients extracted from the null-model data and those obtained from the original connectome data, we quantified the pairwise similarity as follows:

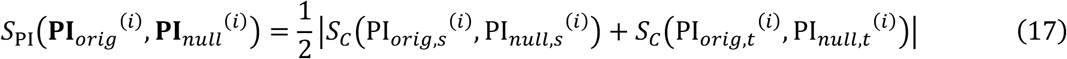

where 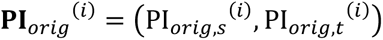 and 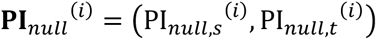 are the *i*th pair of wiring PI vectors extracted from original connectome data and null-model data, respectively, and *S*_*C*_(***x***, ***y***) denotes cosine similarity defined in Eq. (16). A higher value of *S*_9:_ indicates greater similarity between the null model-derived wiring PI and that from the original data.

## Data Availability

The data supporting this study were obtained from the Allen Brain Atlas and supplemented with datasets from previously published articles, as described in the Methods section.

## Code Availability

The SPERRFY is developed under MATLAB 2024. All code used in the current study will be available on the GitHub repository after publication.

## Acknowledgements

This work was supported by JST, the establishment of university fellowships towards the creation of science technology innovation (grant number JPMJFS2129; to J.K.), JST SPRING (grant number JPMJSP2132; to J.K.), Moonshot R&D–MILLENNIA Program (grant number JPMJMS2024-9; to H.N.) and the Cooperative Study Program of Exploratory Research Center on Life and Living Systems (ExCELLS; program number 19–102; to H.N.). We would like to thank the members of Honda Lab and other laboratories of the mathematical biology group at Hiroshima University for their insightful discussion. Additionally, we acknowledge the assistance of ChatGPT in refining the language of the manuscript.

## Author Contributions

H.N. conceived the project. J.K. developed the model and analyzed the data. J.K., Y.Y., R.H., and H.N. wrote the manuscript with input from all the authors.

## Supplementary Figures

**Supplementary Figure 1:**
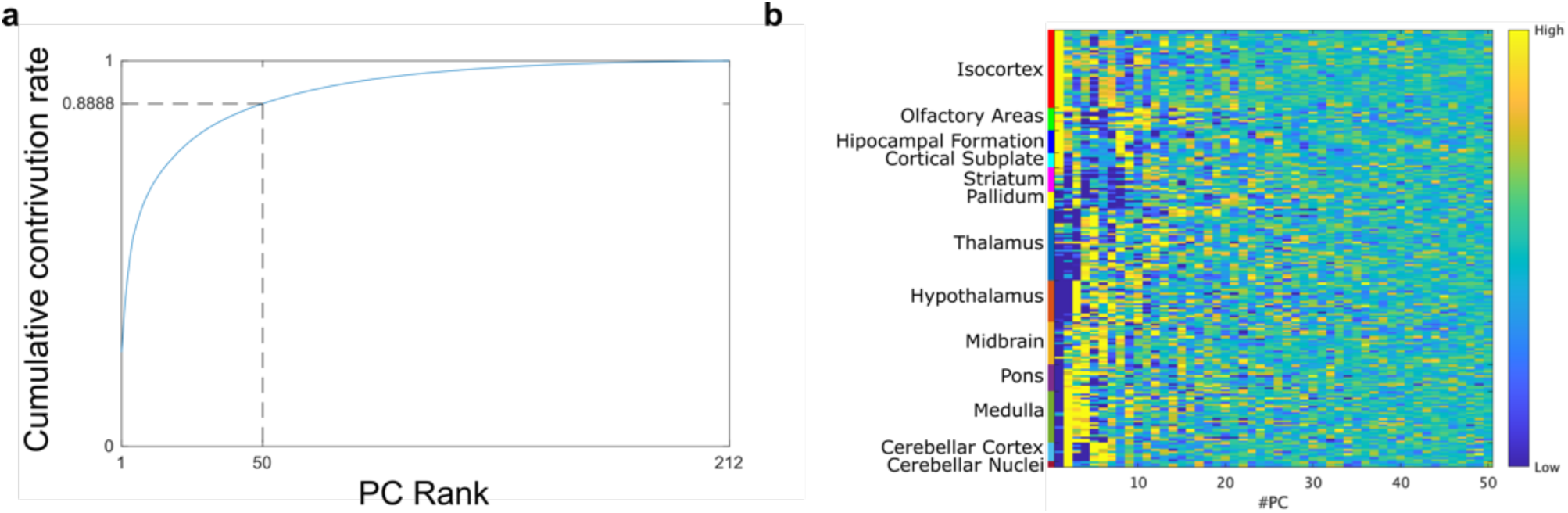
Principal component analysis (PCA) of spatial gene expression data. **a:** Cumulative contribution ratio of the top principal components. The dashed line indicates the 50th principal component, which was used as the dimensionality cutoff in the analysis. **b:** Heatmap of the PCA-reduced gene expression data, showing the expression profiles of 213 brain regions across the top 50 principal components (PCs). Color represents expression values.

**Supplementary Figure 2:**
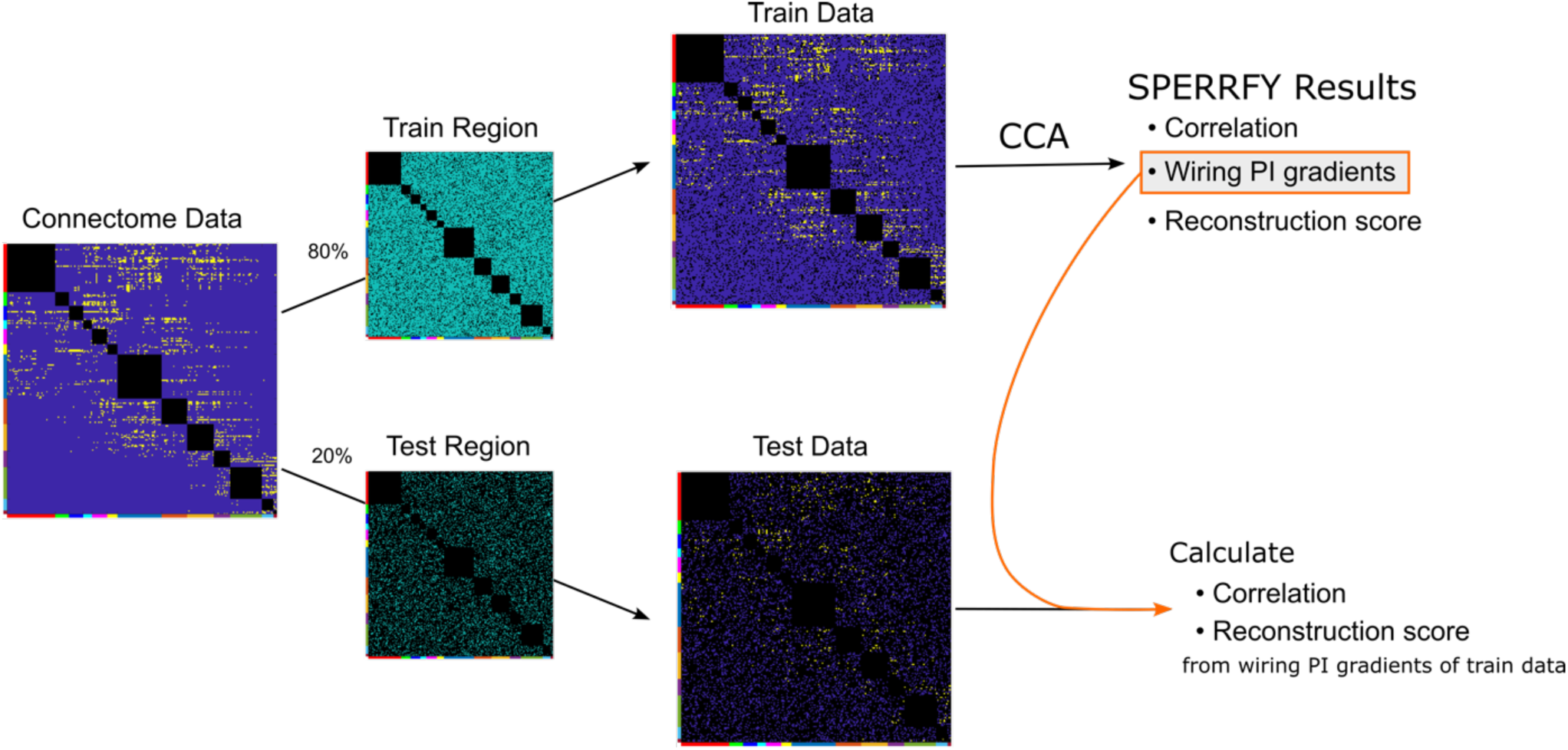
Schematic illustration of the hold-out validation procedure. The original connectome matrix was randomly split into a training region and a test region. CCA was applied only to the training data to extract wiring PI gradients. These gradients were then used to compute correlation coefficients and reconstruction performance on the held-out test data, thereby assessing the generalizability of SPERRFY results.

**Supplementary Figure 3:**
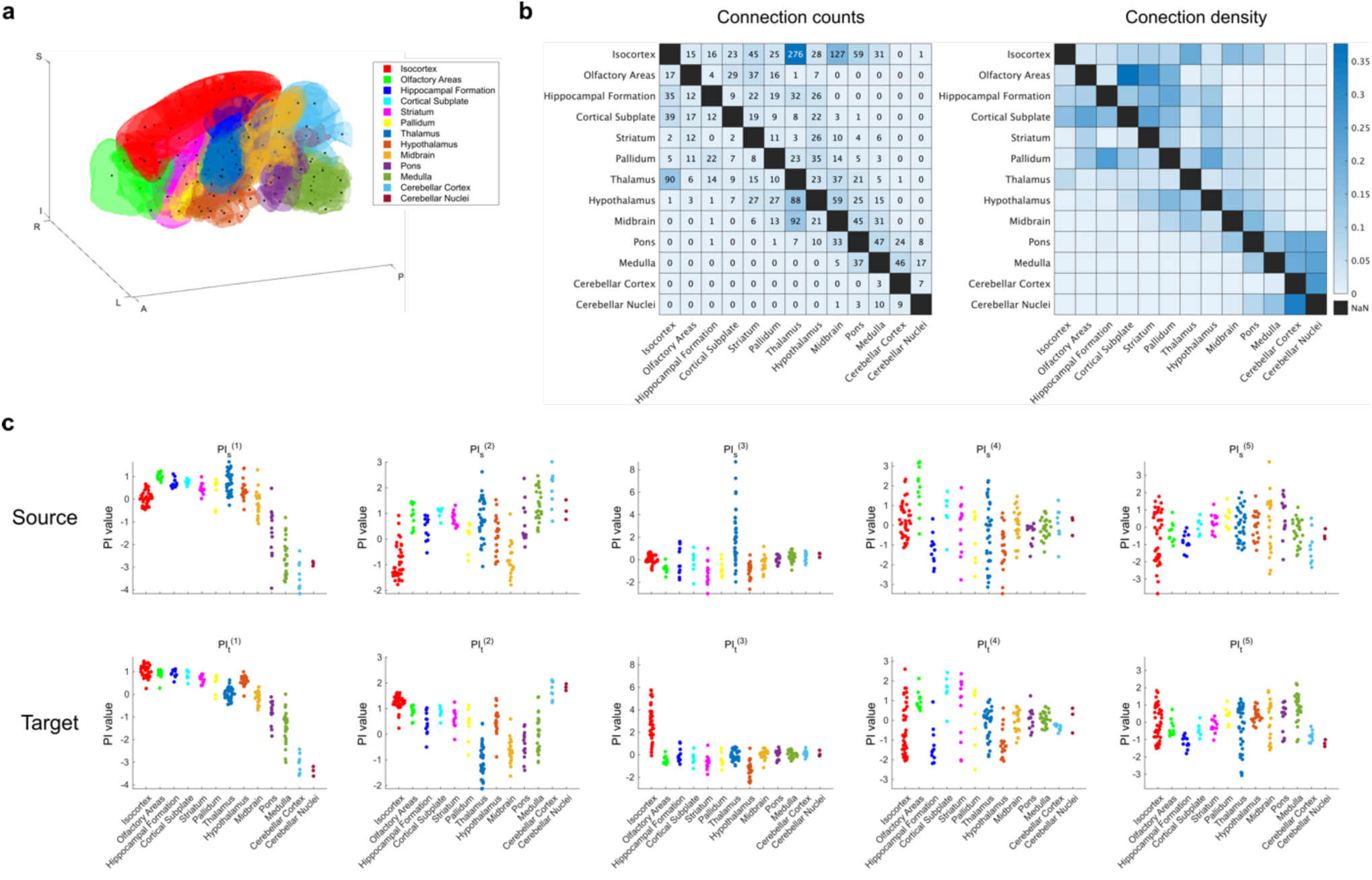
Properties of major brain regions (MRs) used in the analysis. **a:** 3D spatial visualization of the 13 major regions (MRs) in the mouse brain. **b:** (Left) Number of connections between each MR pair. (Right) Connection density normalized by the number of possible connections. **c:** Distribution of wiring PI values across brain regions, grouped by MR. Each dot represents a single brain region. Top: source PI components. Bottom: target PI components.

**Supplementary Figure 4:**
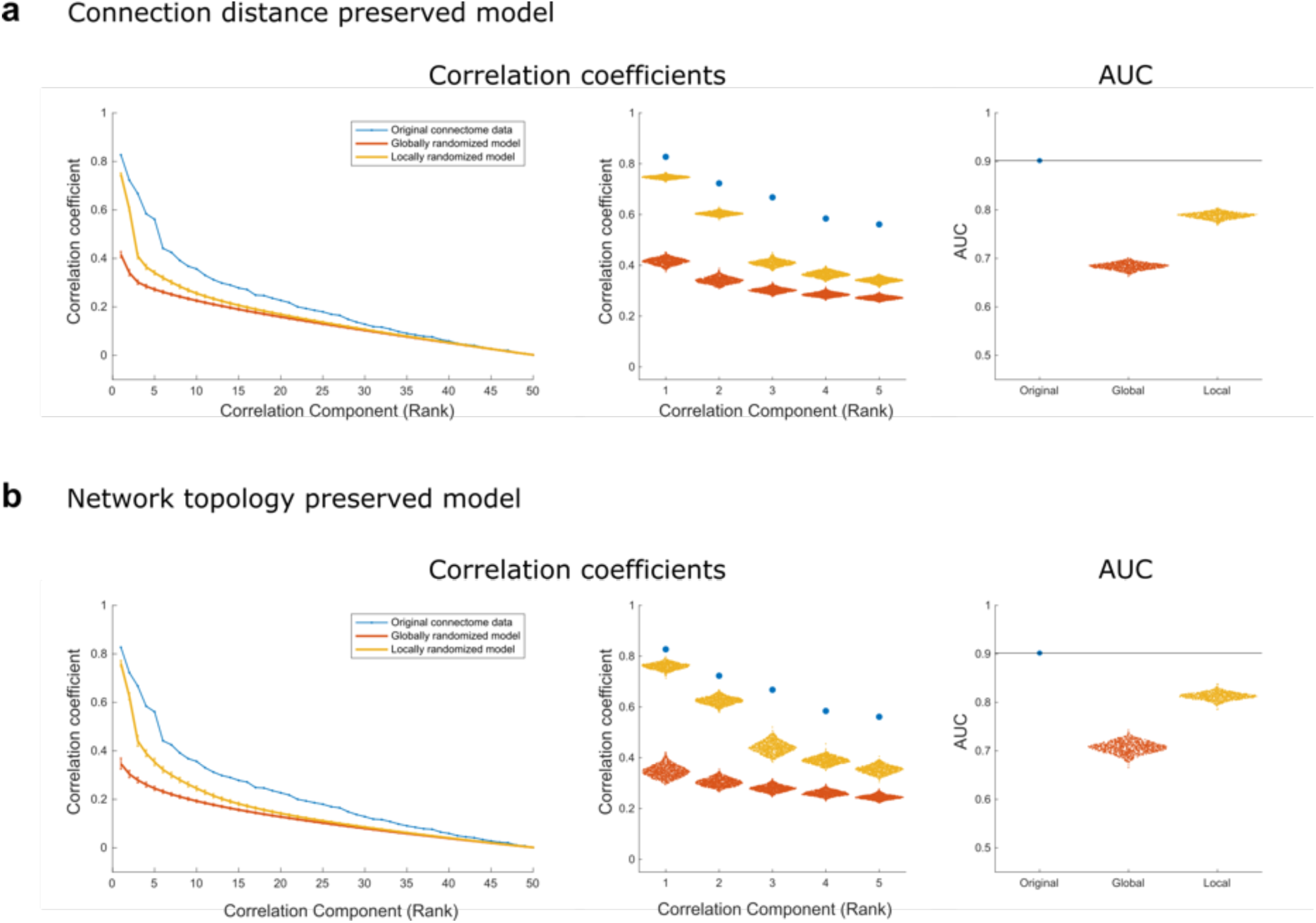
Comparison with additional null models incorporating biological constraints. **a:** Null model in which the distribution of connection distances is preserved to match that of the original connectome. **b:** Null model in which network topology is preserved by shuffling brain region labels while keeping the original connection matrix intact. For both (A) and (B): (Left) – mean correlation coefficients of wiring PI components across ranks (blue: original, red: globally randomized, yellow: constrained randomized). (Middle) – distributions of the top five correlation coefficients. (Right) – distributions of AUC values for connectome reconstruction.

**Supplementary Figure 5:**
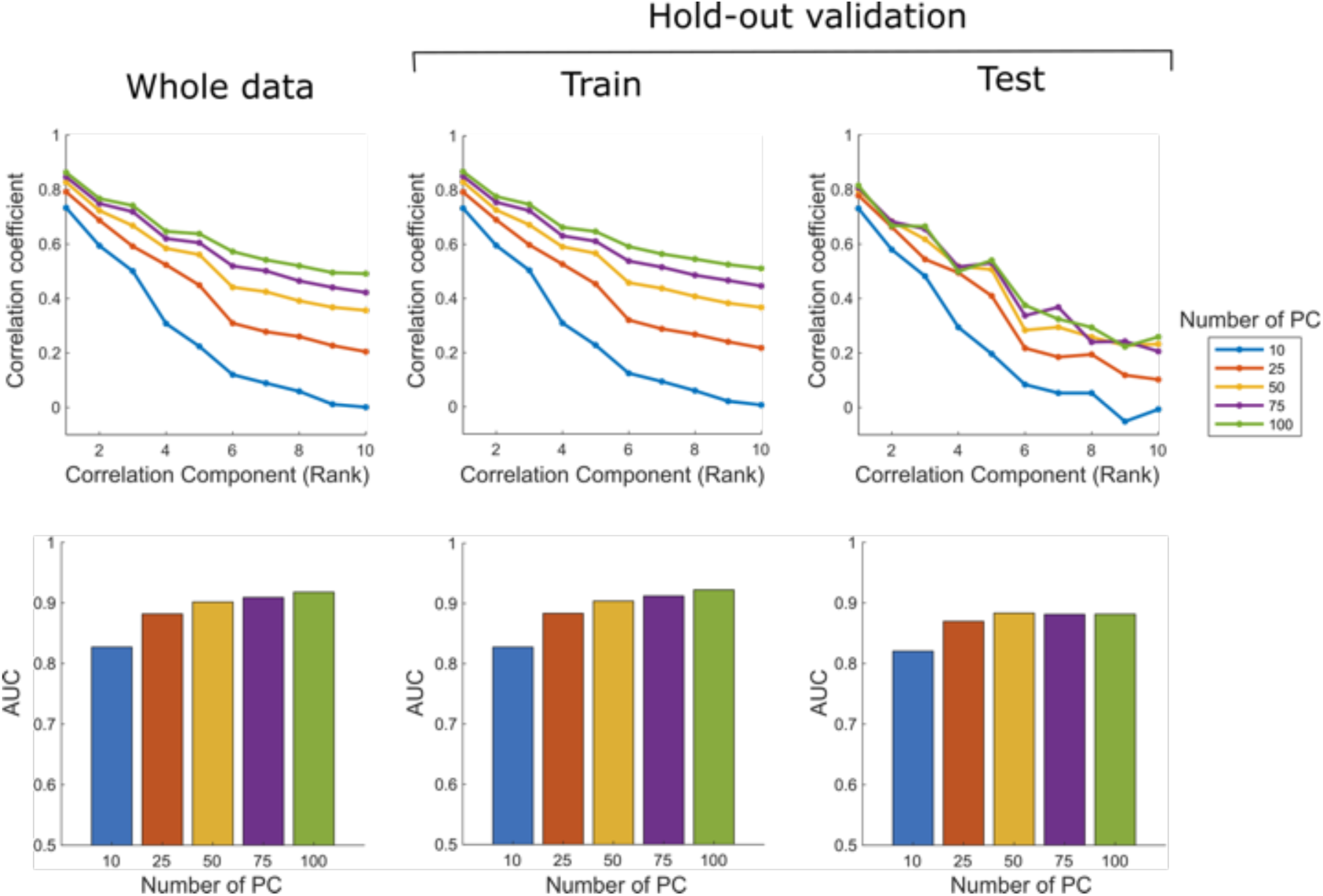
Evaluation of the effect of PCA dimensionality on SPERRFY performance. (Top) Correlation coefficients of the top 10 wiring PI components across different numbers of principal components (PCs) used for gene expression representation (10, 25, 50, 75, 100). (Bottom) AUC scores for connectome reconstruction evaluated on the whole dataset and under hold-out validation (train/test split).

## References

1. Tessier-Lavigne, M. & Goodman, C. S. The Molecular Biology of Axon Guidance. Science 274, 1123–1133 (1996).

2. Dickson, B. J. Molecular Mechanisms of Axon Guidance. Science 298, 1959–1964 (2002).

3. Yu, T. W. & Bargmann, C. I. Dynamic regulation of axon guidance. Nat Neurosci 4, 1169–1176 (2001).

4. Hong, K., Nishiyama, M., Henley, J., Tessier-Lavigne, M. & Poo, M. Calcium signalling in the guidance of nerve growth by netrin-1. Nature 403, 93–98 (2000).

5. Nishiyama, M. et al. Cyclic AMP/GMP-dependent modulation of Ca2+ channels sets the polarity of nerve growth-cone turning. Nature 423, 990–995 (2003).

6. Nakamura, F., Kalb, R. G. & Strittmatter, S. M. Molecular basis of semaphorin-mediated axon guidance. Journal of Neurobiology 44, 219–229 (2000).

7. O’Leary, D. D. M. & McLaughlin, T. Mechanisms of retinotopic map development: Ephs, ephrins, and spontaneous correlated retinal activity. in Progress in Brain Research vol. 147 43–65 (Elsevier, 2005).

8. Scicolone, G., Ortalli, A. L. & Carri, N. G. Key roles of Ephs and ephrins in retinotectal topographic map formation. Brain Research Bulletin 79, 227–247 (2009).

9. Triplett, J. W. & Feldheim, D. A. Eph and ephrin signaling in the formation of topographic maps. Seminars in Cell & Developmental Biology 23, 7–15 (2012).

10. Egea, J. & Klein, R. Bidirectional Eph–ephrin signaling during axon guidance. Trends in Cell Biology 17, 230–238 (2007).

11. Knöll, B. & Drescher, U. Ephrin-As as receptors in topographic projections. Trends in Neurosciences 25, 145–149 (2002).

12. Dufour, A. et al. Area Specificity and Topography of Thalamocortical Projections Are Controlled by ephrin/Eph Genes. Neuron 39, 453–465 (2003).

13. Marín, O., Blanco, M. J. & Nieto, M. A. Differential Expression of Eph Receptors and Ephrins Correlates with the Formation of Topographic Projections in Primary and Secondary Visual Circuits of the Embryonic Chick Forebrain. Developmental Biology 234, 289–303 (2001).

14. Zipursky, S. L. & Sanes, J. R. Chemoaffinity Revisited: Dscams, Protocadherins, and Neural Circuit Assembly. Cell 143, 343–353 (2010).

15. Sperry, R. W. Chemoaffinity in the orderly growth of nerve fiber patterns and connections*. Proceedings of the National Academy of Sciences 50, 703–710 (1963).

16. Cheng, H.-J., Nakamoto, M., Bergemann, A. D. & Flanagan, J. G. Complementary gradients in expression and binding of ELF-1 and Mek4 in development of the topographic retinotectal projection map. Cell 82, 371–381 (1995).

17. Yuasa, J., Hirano, S., Yamagata, M. & Noda, M. Visual projection map specified by topographic expression of transcription factors in the retina. Nature 382, 632–635 (1996).

18. Weth, F., Fiederling, F., Gebhardt, C. & Bastmeyer, M. Chemoaffinity in topographic mapping revisited – Is it more about fiber–fiber than fiber–target interactions? Seminars in Cell & Developmental Biology 35, 126–135 (2014).

19. Naoki, H. Revisiting chemoaffinity theory: Chemotactic implementation of topographic axonal projection. PLOS Computational Biology 13, e1005702 (2017).

20. Scalia, F. Synapse formation in the olfactory cortex by regenerating optic axons: Ultrastructural evidence for polyspecific chemoaffinity. Journal of Comparative Neurology 263, 497–513 (1987).

21. Wang, F., Nemes, A., Mendelsohn, M. & Axel, R. Odorant Receptors Govern the Formation of a Precise Topographic Map. Cell 93, 47–60 (1998).

22. Van Essen, D. C. et al. The WU-Minn Human Connectome Project: An overview. NeuroImage 80, 62–79 (2013).

23. Van Essen, D. C. et al. The Human Connectome Project: A data acquisition perspective. NeuroImage 62, 2222–2231 (2012).

24. Oh, S. W. et al. A mesoscale connectome of the mouse brain. Nature 508, 207–214 (2014).

25. Tian, X. et al. An integrated resource for functional and structural connectivity of the marmoset brain. Nat Commun 13, 7416 (2022).

26. Skibbe, H. et al. The Brain/MINDS Marmoset Connectivity Resource: An open-access platform for cellular-level tracing and tractography in the primate brain. PLOS Biology 21, e3002158 (2023).

27. Scheffer, L. K., et al. A connectome and analysis of the adult Drosophila central brain. eLife 9, e57443 (2020).

28. Dorkenwald, S. et al. FlyWire: online community for whole-brain connectomics. Nat Methods 19, 119–128 (2022).

29. Winding, M. et al. The connectome of an insect brain. Science 379, eadd9330 (2023).

30. Schlegel, P. et al. Whole-brain annotation and multi-connectome cell typing of Drosophila. Nature 634, 139–152 (2024).

31. Shen, E. H., Overly, C. C. & Jones, A. R. The Allen Human Brain Atlas: Comprehensive gene expression mapping of the human brain. Trends in Neurosciences 35, 711–714 (2012).

32. Lein, E. S. et al. Genome-wide atlas of gene expression in the adult mouse brain. Nature 445, 168–176 (2007).

33. Ng, L. et al. An anatomic gene expression atlas of the adult mouse brain. Nat Neurosci 12, 356–362 (2009).

34. Henry, A. M. & Hohmann, J. G. High-resolution gene expression atlases for adult and developing mouse brain and spinal cord. Mamm Genome 23, 539–549 (2012).

35. Thompson, C. L. et al. A High-Resolution Spatiotemporal Atlas of Gene Expression of the Developing Mouse Brain. Neuron 83, 309–323 (2014).

36. Shimogori, T. et al. Digital gene atlas of neonate common marmoset brain. Neuroscience Research 128, 1–13 (2018).

37. Robinson, S. W., Herzyk, P., Dow, J. A. T. & Leader, D. P. FlyAtlas: database of gene expression in the tissues of Drosophila melanogaster. Nucleic Acids Research 41, D744–D750 (2013).

38. French, L. & Pavlidis, P. Relationships between Gene Expression and Brain Wiring in the Adult Rodent Brain. PLOS Computational Biology 7, e1001049 (2011).

39. Fornito, A., Arnatkevičiūtė, A. & Fulcher, B. D. Bridging the Gap between Connectome and Transcriptome. Trends in Cognitive Sciences 23, 34–50 (2019).

40. Fulcher, B. D. & Fornito, A. A transcriptional signature of hub connectivity in the mouse connectome. Proceedings of the National Academy of Sciences 113, 1435–1440 (2016).

41. Mills, B. D. et al. Correlated Gene Expression and Anatomical Communication Support Synchronized Brain Activity in the Mouse Functional Connectome. J. Neurosci. 38, 5774–5787 (2018).

42. Hardoon, D. R., Szedmak, S. & Shawe-Taylor, J. Canonical Correlation Analysis: An Overview with Application to Learning Methods. Neural Computation 16, 2639–2664 (2004).

43. Sunkin, S. M. et al. Allen Brain Atlas: an integrated spatio-temporal portal for exploring the central nervous system. Nucleic Acids Research 41, D996–D1008 (2013).

44. Wang, Q. et al. The Allen Mouse Brain Common Coordinate Framework: A 3D Reference Atlas. Cell 181, 936–953.e20 (2020).

45. Ringnér, M. What is principal component analysis? Nat Biotechnol 26, 303–304 (2008).

46. Yadav, S. & Shukla, S. Analysis of k-Fold Cross-Validation over Hold-Out Validation on Colossal Datasets for Quality Classification. in 2016 IEEE 6th International Conference on Advanced Computing (IACC) 78–83 (2016). doi:10.1109/IACC.2016.25.

47. Pasquale, E. B. Eph-Ephrin Bidirectional Signaling in Physiology and Disease. Cell 133, 38–52 (2008).

48. Scheefhals, N., Westra, M. & MacGillavry, H. D. mGluR5 is transiently confined in perisynaptic nanodomains to shape synaptic function. Nat Commun 14, 1–20 (2023).

49. Loerwald, K. W., Patel, A. B., Huber, K. M. & Gibson, J. R. Postsynaptic mGluR5 promotes evoked AMPAR-mediated synaptic transmission onto neocortical layer 2/3 pyramidal neurons during development. Journal of Neurophysiology 113, 786–795 (2015).

50. Elmasri, M. et al. Synaptic Dysfunction by Mutations in GRIN2B: Influence of Triheteromeric NMDA Receptors on Gain-of-Function and Loss-of-Function Mutant Classification. Brain Sciences 12, 789 (2022).

51. Rodríguez-Palmero, A., et al. *DLG4*-related synaptopathy: a new rare brain disorder. Genetics in Medicine 23, 888–899 (2021).

52. Lisman, J., Yasuda, R. & Raghavachari, S. Mechanisms of CaMKII action in long-term potentiation. Nat Rev Neurosci 13, 169–182 (2012).

53. Chivatakarn, O., Kaneko, S., He, Z., Tessier-Lavigne, M. & Giger, R. J. The Nogo-66 Receptor NgR1 Is Required Only for the Acute Growth Cone-Collapsing But Not the Chronic Growth-Inhibitory Actions of Myelin Inhibitors. J. Neurosci. 27, 7117–7124 (2007).

54. Pancho, A. et al. Modifying PCDH19 levels affects cortical interneuron migration. Front. Neurosci. 16, (2022).

55. Piao, X. et al. G Protein-Coupled Receptor-Dependent Development of Human Frontal Cortex. Science 303, 2033–2036 (2004).

56. Li, S. et al. GPR56 Regulates Pial Basement Membrane Integrity and Cortical Lamination. J. Neurosci. 28, 5817–5826 (2008).

57. Gonda, Y., Namba, T. & Hanashima, C. Beyond Axon Guidance: Roles of Slit-Robo Signaling in Neocortical Formation. Front. Cell Dev. Biol. 8, (2020).

58. Okada, A. et al. Boc is a receptor for sonic hedgehog in the guidance of commissural axons. Nature 444, 369–373 (2006).

59. Ferent, J. et al. Boc Acts via Numb as a Shh-Dependent Endocytic Platform for Ptch1 Internalization and Shh-Mediated Axon Guidance. Neuron 102, 1157–1171.e5 (2019).

60. Kaiser, M. A tutorial in connectome analysis: Topological and spatial features of brain networks. NeuroImage 57, 892–907 (2011).

61. Drager, U. C. & Hubel, D. H. Topography of visual and somatosensory projections to mouse superior colliculus. Journal of Neurophysiology 39, 91–101 (1976).

62. Kaas, J. H. Topographic Maps are Fundamental to Sensory Processing. Brain Research Bulletin 44, 107–112 (1997).

63. Merzenich, M. M. et al. Topographic reorganization of somatosensory cortical areas 3b and 1 in adult monkeys following restricted deafferentation. Neuroscience 8, 33–55 (1983).

64. Beck, P. D., Pospichal, M. W. & Kaas, J. H. Topography, architecture, and connections of somatosensory cortex in opossums: Evidence for five somatosensory areas. Journal of Comparative Neurology 366, 109–133 (1996).

65. Takahashi, Y. K., Kurosaki, M., Hirono, S. & Mori, K. Topographic Representation of Odorant Molecular Features in the Rat Olfactory Bulb. Journal of Neurophysiology 92, 2413–2427 (2004).

66. Gogos, J. A., Osborne, J., Nemes, A., Mendelsohn, M. & Axel, R. Genetic Ablation and Restoration of the Olfactory Topographic Map. Cell 103, 609–620 (2000).

67. Vassar, R. et al. Topographic organization of sensory projections to the olfactory bulb. Cell 79, 981–991 (1994).

68. Luskin, M. B. & Price, J. L. The topographic organization of associational fibers of the olfactory system in the rat, including centrifugal fibers to the olfactory bulb. Journal of Comparative Neurology 216, 264–291 (1983).

69. Imai, T., Sakano, H. & Vosshall, L. B. Topographic Mapping̶The Olfactory System. Cold Spring Harb Perspect Biol 2, a001776 (2010).

70. López-Bendito, G. & Molnár, Z. Thalamocortical development: how are we going to get there? Nat Rev Neurosci 4, 276–289 (2003).

71. Catalano, S. M., Robertson, R. T. & Killackey, H. P. Individual axon morphology and thalamocortical topography in developing rat somatosensory cortex. Journal of Comparative Neurology 367, 36–53 (1996).

72. Molnár, Z., Adams, R. & Blakemore, C. Mechanisms Underlying the Early Establishment of Thalamocortical Connections in the Rat. J. Neurosci. 18, 5723–5745 (1998).

73. Ji, S., Fakhry, A. & Deng, H. Integrative analysis of the connectivity and gene expression atlases in the mouse brain. NeuroImage 84, 245–253 (2014).

74. Fakhry, A. & Ji, S. High-resolution prediction of mouse brain connectivity using gene expression patterns. Methods 73, 71–78 (2015).

75. Sun, S., Torok, J., Mezias, C., Ma, D. & Raj, A. Spatial cell-type enrichment predicts mouse brain connectivity. Cell Reports 42, 113258 (2023).

76. Fulcher, B. D., Arnatkeviciute, A. & Fornito, A. Overcoming false-positive gene-category enrichment in the analysis of spatially resolved transcriptomic brain atlas data. Nat Commun 12, 2669 (2021).

77. Bullmore, E. & Sporns, O. The economy of brain network organization. Nat Rev Neurosci 13, 336–349 (2012).

78. Gao, W. et al. Temporal and Spatial Evolution of Brain Network Topology during the First Two Years of Life. PLOS ONE 6, e25278 (2011).

79. Li, H. et al. Mapping fetal brain development based on automated segmentation and 4D brain atlasing. Brain Struct Funct 226, 1961–1972 (2021).

80. Braun, E. et al. Comprehensive cell atlas of the first-trimester developing human brain. Science 382, eadf1226 (2023).

81. Jiang, F. et al. Simultaneous profiling of spatial gene expression and chromatin accessibility during mouse brain development. Nat Methods 20, 1048–1057 (2023).

82. Sonoda, M. et al. Six-dimensional dynamic tractography atlas of language connectivity in the developing brain. Brain 144, 3340–3354 (2021).

83. Li, Y. et al. Spatiotemporal transcriptome atlas reveals the regional specification of the developing human brain. Cell 186, 5892–5909.e22 (2023).

